# An essential mycolate remodeling program for mycobacterial adaptation in host cells

**DOI:** 10.1101/354431

**Authors:** Eliza J.R. Peterson, Rebeca Bailo, Alissa C. Rothchild, Mario Arrieta-Ortiz, Amardeep Kaur, Min Pan, Dat Mai, Charlotte Cooper, Alan Aderem, Apoorva Bhatt, Nitin S. Baliga

## Abstract

The success of *Mycobacterium tuberculosis* (MTB) stems from its ability to remain hidden from the immune system within macrophages. Here, we report a new technology (Path-seq) to sequence miniscule amounts of MTB transcripts within up to million-fold excess host RNA. Using Path-seq we have discovered a novel transcriptional program for *in vivo* mycobacterial cell wall remodeling when the pathogen infects alveolar macrophages in mice. We have discovered that MadR transcriptionally modulates two mycolic acid desaturases *desA1/A2* to initially promote cell wall remodeling upon *in vitro* macrophage infection and, subsequently, reduces mycolate biosynthesis upon entering dormancy. We demonstrate that disrupting MadR program is lethal to diverse mycobacteria making this evolutionarily conserved regulator a prime antitubercular target for both early and late stages of infection.

**One Sentence Summary:** Novel technology (Path-seq) discovers cell wall remodeling program during *Mycobacterium tuberculosis* infection of macrophages

## Main Text

*Mycobacterium tuberculosis* (MTB) infection occurs by inhalation of bacilli-containing aerosols. Alveolar macrophages, which line the airway, are the first host cells to phagocytize the bacteria. This initial contact of MTB with alveolar macrophages begins a complex battle between bacterial virulence and host immunity, orchestrated in large part by intricate gene regulatory pathways(*1*, *2*). As such, measuring gene expression *in vivo* is central to our understanding of TB disease control and progression(*3*).

RNA-seq provides a sensitive method for global gene expression analysis. Specific for infection biology, dual RNA-seq methods have allowed simultaneous profiling of host and pathogen RNA. However, the striking excess of eukaryotic over bacteria RNA limits the coverage of pathogen transcripts in dual RNA-seq studies(*4*–*8*), and methods to partially enrich for bacterial transcripts have had limited success(*9*, *10*). It is clear more sensitive approaches are needed to profile the transcriptional state of the pathogen during infection, especially *in vivo*.

To improve the coverage of pathogen transcripts, we made use of biotinylated oligonucleotide baits that are complementary to the pathogen transcriptome. The baits are hybridized to mixed host-pathogen RNA and used to enrich pathogen transcripts for sequencing. We applied our pathogen-sequencing (Path-seq) method to explore transcriptional changes in MTB following infection in mice. Here Path-seq has led to discovery that MTB transcriptionally regulate mycolic acids during infection of host cells, influencing virulence and persistence of the pathogen.

## RESULTS and DISCUSSION

### Development of Path-seq

To enrich the bacterial pathogen transcripts, we used Agilent eArray(*11*) to create a custom bait library that covers all MTB transcripts at even intervals. Our MTB library contains 35,624 probes, each with biotinylated oligonucleotides of 120 base lengths. The bait library composition is modular and can be designed to cover specific transcripts of interest. Similarly, transcripts such as rRNA can be excluded or gene sequences altered for polymorphisms found in clinical strains(*12*). For this study, we chose all transcripts of MTB H37Rv for complete coverage and comparison to standard RNA-seq results.

To assess enrichment of pathogen transcripts, we first used RNA isolated from murine bone marrow derived marcrophages (BMDMs) spiked with 0.1% MTB RNA. A typical mammalian cell contains on the order of 20 picograms of RNA, which is roughly two orders of magnitude more than a single bacterial cell(*13*). Accounting for BMDMs that might not be infected and based on intracellular sequencing studies from literature(*4*, *8*), we estimated 0.1% pathogen RNA would be representative of a typical *in vitro* infection. We performed double rRNA depletion using Illumina Ribo-Zero Gold Epidemiology Kit and used the SureSelect protocol to generate strand-specific libraries for sequencing. Half of the library was then indexed for sequencing as the “RNA-seq” sample and the other was hybridized to the probes, amplified and indexed as the “Path-seq” sample (**Fig. 1A**). We performed three replicate experiments of the mock infection using the same MTB RNA. With the probe hybridization, the percentages of reads aligned to MTB were increased up to 840-fold. Both the normalized read counts (**Fig. 1B**) and enrichment efficiency (inset **Fig. 1B**) were highly reproducible across three replicate samples. Repeating the Path-seq method with spiked RNA samples, we increased the proportion of macrophage RNA and were able to quantify MTB transcripts from one millionth of the host RNA (1.75% of all reads aligned to MTB genomes).

**Fig. 1.**
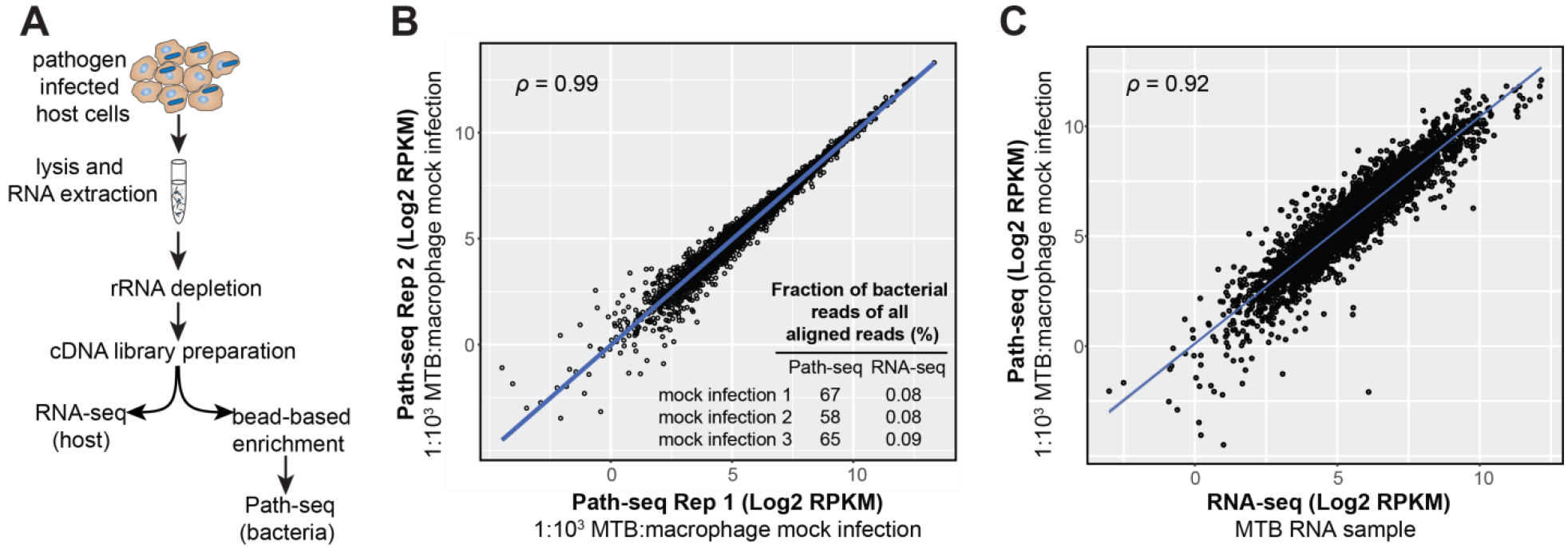
Path-seq workflow and validation. (**A**) Total RNA is extracted from infected cells, depleted of rRNA, and cDNA libraries prepared. Libraries can then be indexed and sequenced directly for host transcripts or enriched using pathogen specific oligonucleotides bound to beads. After hybridization, enriched libraries are indexed, sequenced and reads assigned to host or pathogen genomes *in silico*. (**B**) Correlation between replicate mock infections. Path-seq reads were recovered from samples of macrophage RNA spiked with 0.1% MTB RNA. Scatter plot of log2 RPKM values is shown with Pearson correlation, *P*-value < 0.0001. Inset summarizes the mock infection replicates and their fraction of MTB reads (of all aligned reads) from Path-seq and standard RNA-seq methods. (**C**) Correlation between MTB RNA sequenced by RNA-seq (Illumina TruSeq library prep) and the same MTB RNA sample combined with macrophage RNA at 1:1000 ratio and processed using Path-seq method. Scatter plot of log2 RPKM values is shown with Pearson correlation, *P*-value < 0.0001.

To validate the enrichment protocol yielded quantitatively reliable read counts, we investigated the correlation between the RPKMs obtained from the sequencing of RNA from *in vitro* grown MTB, without enrichment (RNA-seq), and the RPKM values obtained with the enrichment protocol (Path-seq) using the same 0.1% MTB RNA with host RNA (BMDMs). Even using different library preparation kits (Illumina for RNA-seq and Agilent for Path-seq), the correlation of RPKMs was 0.92-0.93 (**Fig. 1C**), demonstrating the enrichment process was efficient and accurate for gene expression analysis.

### Analysis of MTB transcriptome during *in vivo* infection using Path-seq method

Little is known about the transcriptional state of the pathogen during infection of animal models(*14*); technical challenges have limited these studies. Given the enrichment capabilities of the Path-seq method, we evaluated the use of the approach to study the transcriptome of MTB isolated from alveolar macrophages (AMs) of infected mice. Using fluorescence-activated cell sorting (FACS) and gating strategies to isolate AMs from bronchoalveolar lavage (BAL) of mice infected with mEmerald-expressing MTB, we determined 2% of AMs were infected with MTB at 24 h post infection (data not shown). We isolated AMs from BAL, instead of whole lung tissue, to avoid the harsh digestion step at 37 °C that can alter the transcriptional state of the cells. However, the 2% infected AM population from BAL was limiting for successful RNA extraction. Therefore, we isolated the entire AM population (i.e., infected and bystanders), relying on the Path-seq method to enrich the MTB transcripts, all of which come from intracellular bacteria. RNA extracted from ~2×10^5^ AMs (infected and bystanders) in BAL of 10 mice yielded ~100 μg of total RNA. Therefore, we first evaluated the Path-seq method using 0.3 μg of BMDM RNA spiked with 0.005% MTB RNA, as an estimate of an *in vivo* AM infection. We sequenced two replicates and alignment analysis revealed the percentages of reads that aligned to MTB were 38% and 27%, an approximate 1,000-fold enrichment.

After evaluating the Path-seq methods feasibility for *in vivo* MTB transcriptome analysis, we used flow cytometry to isolate ~6×10^5^ AMs (average of 4.3% of all cells and 83.1% of live, CD45+ cells) in BAL of mice 24 h after infection with wild-type MTB (**Fig. S1A**). Infection, FACS-sorting, and RNA extraction was repeated with three animal groups (**Fig. S1B**), yielding an average ~300 ug total RNA from 30 pooled mice. The Path-seq enrichment was performed and resulted in 17%, 8% and 5% of the entire reads aligning to MTB. We compared the MTB read counts between the *in vivo* samples and extracellular samples grown in media for 24 h, also processed by Path-seq. While the percentage of non-zero reads and total read counts are lower in the *in vivo* samples, the mean count per gene and noise are the same between the two conditions (**Table S1**). This gives us confidence that for genes with detectable reads, we are measuring real expression levels. For differential expression analysis, we excluded any genes with zero counts in all replicates, resulting in 3,505 genes (62%) for comparison between *in vivo* and extracellular MTB. We also analyzed a reduced gene set that excluded genes with zero counts in any replicate, resulting in 1057 genes (20%). Differential expression analysis between *in vivo* intracellular MTB and extracellular MTB, identified 431 differentially expressed genes (log2 fold change < −1.0 or > 1.0 and multiple hypothesis corrected *P*-value < 0.05) from the more stringent gene set (**Data S1**).

Among the most upregulated genes *in vivo* compared to control, were genes encoding aerobic fatty acid desaturases, *desA1* (*Rv0824c*) and *desA2* (*Rv1094*) with log2 fold change of 4.0 (multiple hypothesis corrected *P*-value = 3.0×10^−4^) and 4.7 (adjusted *P*-value = 2.3×10^−4^), respectively. Transposon mutant screens have determined the essentiality of DesA1 and DesA2 for *in vitro* growth of MTB(*15*, *16*). Moreover, we recently demonstrated the essentiality of the *desA1* homolog (*MSMEG_5773*) in *Mycobacterium smegmatis* (MSM)(*17*). We showed that depletion of DesA1 in a conditional mutant caused loss of mycolic acid biosynthesis, reduced cell viability, and accumulation of mono-unsaturated mycolic-acid species consistent with DesA1’s role in the desaturation of mycolic acids(*17*). Crystallography studies were unable to obtain soluble DesA1 but revealed that DesA2 is structurally related to plant fatty acid desaturases and that unordered properties of the protein indicate a specialized role for DesA2(*18*). More biochemical characterization of the desaturases is needed, but the significant up-regulation of *desA1/A2* following *in vivo* infection was interesting and deserved further investigation of their transcriptional control.

### Genome-wide expression analysis during *in vitro* macrophage infection using Path-seq

Several genome-wide expression studies of MTB challenged with various stresses, such as nutrient starvation(*19*), hypoxia(*20*), and during *in vitro* infection(*21*), have shown that genes involved in mycolic acid biosynthesis are generally downregulated. Granted, none of these studies specifically addressed the regulation of the mycolic acid desaturases, but this opposes what we observed in the *in vivo* infection data. Therefore, we used the Path-seq method to study the dynamics of *desA1/A2* expression at higher resolution following MTB infection of bone marrow derived macrophages (BMDMs). We isolated BMDMs and infected them with MTB at a MOI of 10. Infected cells were collected at 2, 8 and 24 h after infection along with extracellular MTB grown in standard media as control. Total RNA was extracted, depleted of rRNA and handled as described above (**Fig. 1A**). All extracellular MTB samples were processed by Path-seq as well. For the infection samples, we again split each sample into RNA-seq and Path-seq fractions to evaluate the enrichment efficiency and to simultaneously obtain both host and pathogen transcriptomes (**Fig. S2**). We evaluated the percentage of reads that aligned to MTB, and found a consistent 100-fold increase in the enriched vs nonenriched samples across replicates and time points (**Table 1**). With an average 11 million (M) mapped reads for both intracellular (average 13.4 M) and extracellular (average 8.8 M) MTB, we obtained >100x coverage and 5,622 unique features (including ncRNA, UTRs, etc.).

**Table 1.** Mapping statistics from Path-seq or RNA-seq samples of MTB infected BMDMs.

|  | Fraction of MTB reads of<br>all aligned reads (%) |  |
| --- | --- | --- |
|  | Path-seq | RNA-seq |
| Intracellular 2h a | 77.0 | 0.73 |
| Intracellular 2h b | 71.1 | 0.53 |
| Intracellular 2h c | 88.5 | 0.65 |
| Intracellular 8h a | 47.7 | 0.54 |
| Intracellular 8h b | 87.5 | 0.66 |
| Intracellular 8h c | 82.8 | 0.38 |
| Intracellular 24h a | 97.4 | 3.97 |
| Intracellular 24h b | 75.5 | 1.88 |
| Intracellular 24h c | 97.5 | 3.10 |

Raw read counts from intracellular and extracellular MTB (all from Path-seq analysis) were used to generate a tSNE network. This resulted in a network map with extracellular samples clustered closely together, distinct from the intracellular samples and according to their time post infection (**Fig. S3A**). Biological replicates fell into related groups and demonstrated strong correlation in pairwise comparison of RPKMs (**Fig. S3B**). Differential expression of intracellular MTB was calculated relative to extracellular, at each time point using DESeq2. Overall, there were 746, 945, and 412 significant differentially expressed (log2 fold change < −1.0 or > 1.0 and multiple hypothesis adjusted P-value < 0. 01) transcripts at the 2, 8, and 24 h post infection time points (**Data S2**). The most upregulated genes at all time points included genes such as *icl1*, *Rv1129c*, *prpD*, *prpC*, and *fadD19*. The induced expression of these genes is consistent with known alteration in lipid metabolism during infection, enhanced activity of the methylcitrate cycle(*22*), and genetic evidence that MTB utilizes cholesterol from the host during infection(*23*). These carbon metabolizing genes were also found to be up-regulated in microarray analysis of MTB infected BMDMs(*21*), along with a significant overlap of other differentially expressed genes between the datasets (multiple hypothesis corrected *P*-value = 6.6×10^−24^ at 2 h and adjusted *P*-value = 1.5×10^−12^ at 24 h). These data demonstrate that the Path-seq method yielded data consistent with published transcriptional studies of *in vitro* infected host cells. Importantly, the Path-seq method allows for simultaneous expression analysis of host transcripts and additional pathogen features that are not possible in microarray studies.

### *desA1* and *desA2* are induced early during *in vitro* macrophage infection

Interestingly, *desA1* and *desA2* were transiently up-regulated at 2 h following MTB infection of BMDMs, followed by return to levels similar to extracellular MTB at 8 h and 24 h (**Fig. 2A**). This is earlier than the Path-seq data from AMs, which showed induced expression of the desaturases at 24h after *in vivo* infection. In fact, we compared all significantly differentially expressed genes between the two infection models at 24 h and found only a small subset of common genes (59 genes), most of which were up-regulated in both models (**Fig. S4**). Furthermore, tSNE analysis of both *in vitro* and *in vivo* infection experiments clusters the AM samples with the controls, separate from the BMDM infected samples (**Fig. 2B**). We hypothesize that the strikingly different gene expression profiles between the experimental infection models reflects heterogeneity in host cells. Huang *et al* recently demonstrated in mice that MTB has lower bacterial stress in AMs compared to interstitial macrophages at two weeks post infection(*24*). The authors theorize that different host macrophage lineages represent different intracellular environments that are permissive (alveolar macrophages) or restrictive (interstitial macrophages) for MTB growth(*24*). Our data also indicate a difference in the MTB transcriptional state from macrophages of different lineages.

**Fig. 2.**
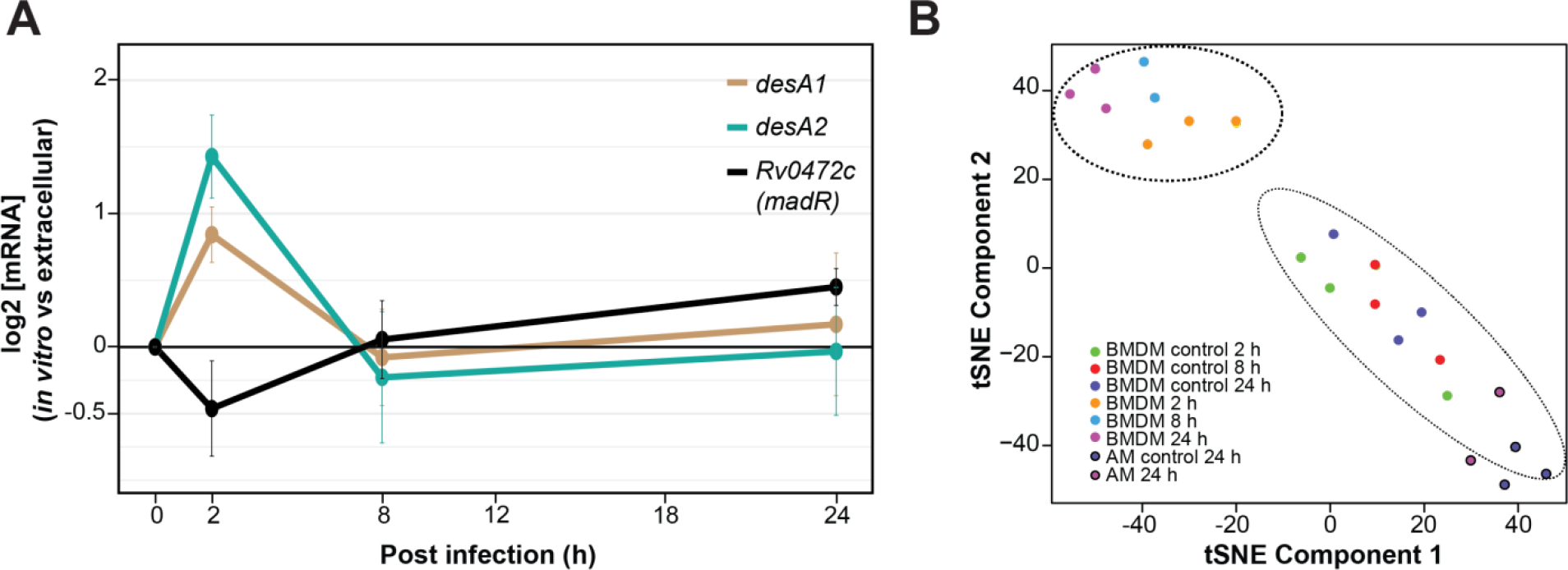
Intracellular *desA1* and *desA2* expression levels from Path-seq of MTB infected mouse AMs (*in vivo*) and BMDMs (*in vitro*). (**A**) Path-seq profiles of *desA1*, *desA2* and Rv0472c in MTB infected BMDMs. Error bars show the standard deviation from three biological samples. Representative results from two experiments are presented. (**B**) tSNE analysis of Path-seq data from both infection models.

### Systems-level comparison of active MTB regulatory networks illustrates differences between infection models

Given the temporal differences in *desA1* and *desA2* expression between the infection models, we employed a computational framework to characterize, at systems-scale, the transcriptional differences between the extracellular and intracellular states for each model. Based on the NetSurgeon algorithm(*25*), we evaluated the role of each TF in the observed gene expression changes given a signed transcriptional network. We constructed a transcriptional network based on ChIP-seq data from overexpression of 178 of 214 TFs in MTB(*26*). Activating and repressing influences of TFs were inferred from consequence of TF overexpression on downstream genes(*27*). Using a data driven transcriptional network of 4,635 interactions, each TF-target gene interaction was weighted according to the multiple hypothesis adjusted P-value from differential expression analysis between intracellular and extracellular conditions. We calculated a relative score for each TF in conditions simulating deletion or overexpression of the TF. These simulations prioritized TF activities (low or high) yielding a transcriptome most similar to the infected state, compared to the control (see **Methods** and summary schematic in **Fig. 3A**). We performed this analysis for each time point and infection model to identify highly ranked TFs (**Fig. 3B**).

**Fig. 3.**
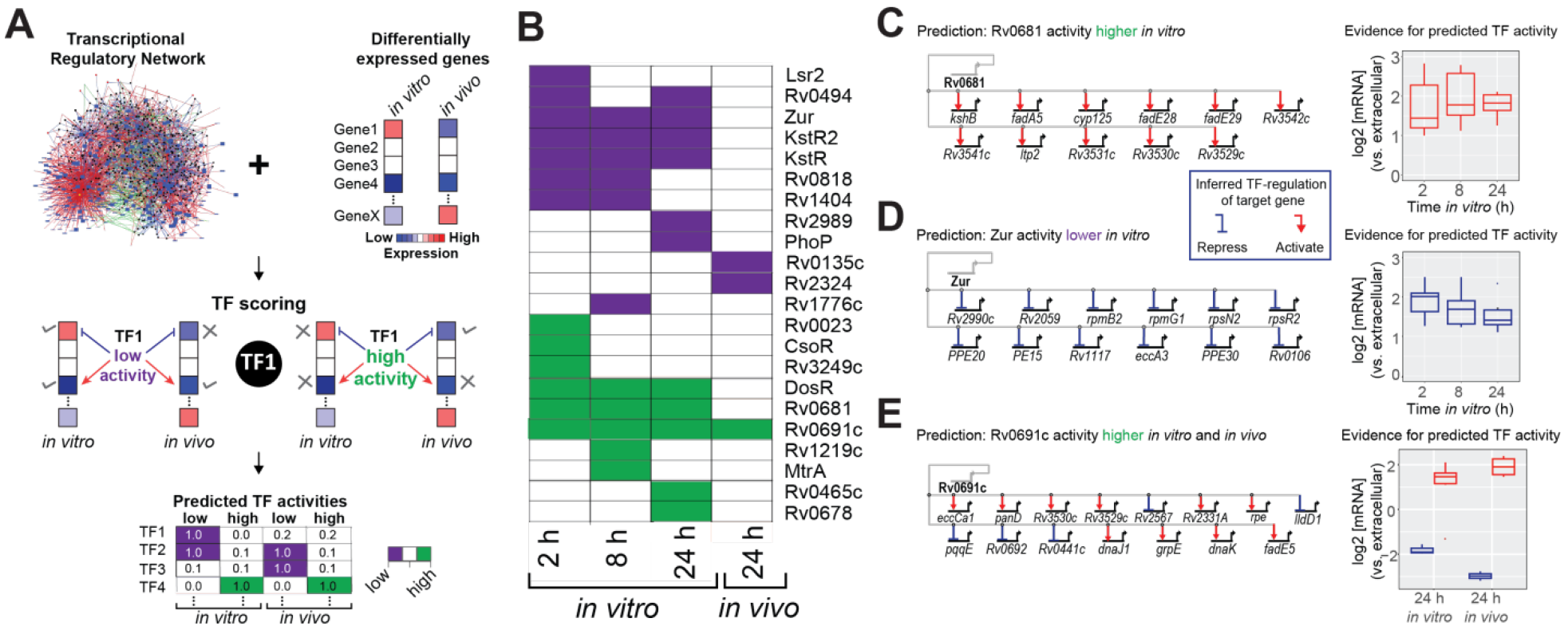
Systems approach to identify active intracellular regulatory networks. (**A**) Schematic of network analysis to identify TFs with activity (high or low) in controlling the transcriptional state of MTB during infection of host cells. (**B**) Heatmap of TFs with low (purple) or high (green) activity at specific time points during *in vitro* or *in vivo* infection. (**C**) Rv0681 regulon genes differentially expressed *in vitro* and the evidence for predicted high Rv0681 activity (induced expression of target genes). (**D**) Zur regulon genes differentially expressed *in vitro* and the evidence for predicted low Zur activity (de-represses target genes). (**E**) Rv0691c regulon genes differentially expressed *in vitro* and *in vivo*; evidence for predicted high Rv0691c activity (increased up- and down-regulation of target genes) at 24 h from both *in vitro* and *in vivo* infection.

From the *in vitro* macrophage infection, many of the TFs had distinct temporal activity, while others were highly ranked across all time points (**Fig. 3B**). These sustained regulons include DosR, which is known to contain a set of ~50 genes that are induced in response to multiple signals including hypoxia, nitrosative stress, and carbon monoxide(*28*–*31*). While DosR regulon induction is typically associated with hypoxic conditions and reactive nitrogen intermediates (RNIs), we observed activation as early as 2 h post infection. Encouragingly, this 2 h induction was also found in the microarray study of MTB infected macrophages, where high DosR regulon expression was sustained until a striking down-regulation at Day 8(*32*). This indicates that the known cues of this regulatory network are present almost immediately during *in vitro* infection. In addition to DosR, two other TFs had high activity across all time points, Rv0681 (**Fig. 3C**) and Rv0691c. Interestingly, both are TetR family transcriptional regulators and conserved across all mycobacterial genomes(*33*), including the drastically reduced *Mycobacterium leprae*. The function of these transcriptional networks is unknown, but suggests their activity is important for survival in both environmental and intracellular niches.

Among the TFs with low predicted activity, KstR and KstR2 were found across all time points of the *in vitro* infection and are known to repress genes required for cholesterol utilization(*34*). Our analysis indicates that deletion of their repressive activity, and increased expression of their target genes, is important for driving the *in vitro* intracellular transcriptional state. This is consistent with the highly expressed cholesterol utilization and methyl citrate cycle genes that we and others have observed(*21*, *35*, *36*). Moreover, this emphasizes the importance of altered carbon metabolism and utilization of host-derived nutrients as key to MTB *in vitro* intracellular adaptation. Another repressor, Zur (previously FurB), had low predicted activity across all time points (**Fig. 3D**). Zur downregulates genes involved in zinc transport(*37*). During MTB infection, macrophages overload the phagosome with copper and zinc as a strategy to poison the pathogen(*38*). However, through multi-faceted resistance mechanisms we do not fully appreciate, MTB is able to protect itself against metal toxicity. Our analysis proposes that reduced Zur activity, results in increased expression of zinc transport genes which could help with regulating zinc levels in MTB during *in vitro* macrophage infection. Interestingly, other regulators of metal content (TFs, uptake and export) were recently found to be required for *in vitro* intracellular growth by high-content imaging of an MTB transposon mutant library(*39*). Leveraging our Path-seq data, we developed a systems-level approach that recapitulates known *in vitro* intracellular regulatory networks and prioritizes others for further experimental testing.

We also applied the same analysis to the *in vivo* expression data (using differentially expressed genes in **Data S1**) to identify transcriptional networks involved in MTBs response to animal infection. Interestingly, we observed very few networks that were active in both infection models. Only Rv0691c was highly ranked at 24 h from both AMs and BMDMs (**Fig. 3E**). In our regulatory network, Rv0691c has ~50 target genes, a subset of which are up- and down-regulated during *in vitro* and *in vivo* infection. The genes in the regulon do not categorize into a certain pathway, but our unbiased analysis suggests the Rv0691c regulon deserves further study for its role in establishing MTB infection both *in vitro* and *in vivo*.

### Identification of *desA1* and *desA2* transcriptional regulator, Rv0472c, and conserved regulation in *M. smegmatis*

Our systems analysis revealed novel and infection-specific regulatory networks. However, none of the identified regulons included *desA1* or *desA2*. We believe this to be a result of regulon size threshold that we implemented to reduce false positives (TFs with at least five targets were considered in this analysis). Therefore, we used the <u>e</u>nvironment and <u>g</u>ene <u>r</u>egulatory <u>i</u>nfluence <u>n</u>etwork (EGRIN)(*40*, *41*) model to discover regulatory mechanisms controlling the expression of the fatty acid desaturases, which were significantly up-regulated following MTB infection of mice. The full description of the algorithms used to construct the EGRIN model are beyond the scope of this work; readers are encouraged to refer to the original paper for more detail(*42*). Briefly, the EGRIN model was constructed through semi-supervised biclustering of a compendium of 2,325 transcriptomes assayed during MTB response to diverse environmental challenges in 53 studies conducted by over 30 different laboratories, guided by biologically informative priors and *de novo* cis-regulatory GRE detection for module assignment(*42*). We demonstrated that the MTB EGRIN model accurately predicts regulatory interactions in a network of 240 modules and validated the model with the DNA binding sites and transcriptional targets from overexpressing >150 MTB transcription factors (TFs)(*26*, *27*). Overall, the EGRIN model is sufficiently predictive to formulate hypotheses of MTB regulatory interactions that respond to various environmental conditions. From the EGRIN model, we identified both desaturases as belonging to module 276 and with predicted regulation by Rv0472c (**Fig. 4A**). Module 276 also contains other genes associated with PDIM biosynthesis and transport. Furthermore, the expression of module genes were found to be significantly correlated with Rv0472c expression under conditions related to oxidative stress and re-aeration(*40*). Rv0472c is a TetR-type TF with homology across all mycobacteria, including *M. leprae*(*33*). When overexpressed in MTB, the TF led to significant repression of 15 genes, but only *desA1* and *desA2* had significant binding of Rv0472c in their promoter region from ChIP-seq analysis(*26*, *27*).

**Fig. 4.**
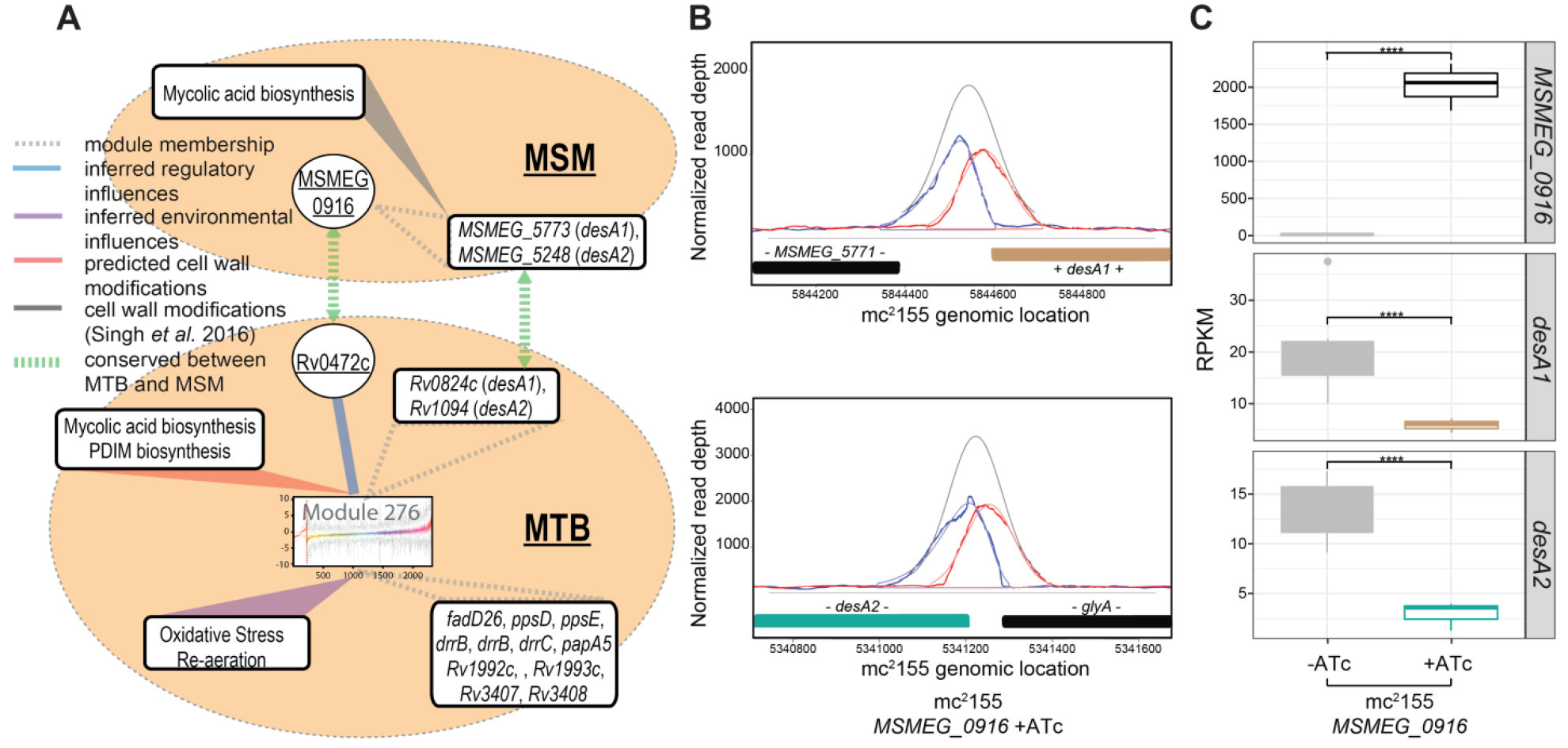
Regulation of *desA1* and *desA2* by Rv0472c (*MSMEG_0916*). (**A**) Regulatory network model of MTB predicts co-regulation of *desA1* and *desA2* and their transcriptional control by Rv0472c. Graphic representation of linkages between Module 276 genes, regulatory and environmental influences, cell wall modifications, and homology to MSM. (**B**) Plot of read pile-ups from MSM with inducible overexpression of *MSMEG_0916* shows ChIP-binding in the promoters of *desA1* and *desA2*. (**C**) Boxplots representing RPKM values from RNA-seq of MSM with inducible overexpression of *MSMEG_0916*. Significant log2 fold change (FC) between uninduced (-ATc) and induced (+ATc) samples for *MSMEG_09l6* (log2 FC = 4.99), *desA1* (log2 FC = −1.32) and *desA2* (log2 FC = −1.72) with multiple hypothesis adjusted *P*-values < 0.0001. Data is from three biological samples.

Given the conservation of Rv0472c across mycobacteria, we hypothesized that overexpression of the MSM homolog, should also repress the desaturases in MSM (**Fig. 4A**). We cloned *MSMEG_0916* into an anhydrotetracycline (ATc)-inducible Gateway shuttle vector as previously described for MTB(*20*, *26*), and transformed into MSM. We induced expression of *MSMEG_0916* for 4h and harvested chromatin samples for ChIP-seq as well as RNA for transcriptional profiling by RNA-seq. Overexpression of *MSMEG_0916* resulted in 9 significant ChIP peaks (*P*-value < 0.01) with a peak score higher than 0.7, as analyzed by DuffyNGS ChIP peak calling method (see **Methods**). Among these, were peaks located in the promoter of the MSM *desA1* and *desA2* (**Fig. 4B**). Additionally, *MSMEG_0916* overexpression resulted in significant repression of *desA1* and *desA2*, with a log2 fold change of −1.32 and −1.72, respectively, compared to uninduced (**Fig. 4C**). The DNA consensus motifs, generated using MEME and DNA-binding data from ChIP-seq, also had significant alignment between Rv0472c and *MSMEG_0916* (**Fig. S5**). The conserved consensus motif is particularly interesting, given that *desA1* and *desA2* are the only regulatory targets shared by the TFs.

### Inducible overexpression of *MSMEG_0916* or *Rv0472c* causes loss of mycobacterial viability and reduction in mycolate biosynthesis

Motivated by our identification of Rv0472c and *MSMEG_0916* as controlling the expression of *desA1* and *desA2*, we hypothesized that overexpression-mediated repression of the desaturases should also have phenotypes similar to the *desA1* knockout that we previously characterized (reduced viability following loss of mycolate biosynthesis)(*17*). We tested the viability of the TF overexpression strains by spotting serial ten-fold dilutions of cultures on agar plates with or without ATc. Plates with *MSMEG_0916* were incubated for 1 week and growth patterns indicated that the presence of ATc resulted in a 2-log fold reduction in CFU counts (**Fig. 5A**). In comparison, plates containing the parental MSM strain showed no change in CFUs with the presence or absence of ATc (**Fig. 5A**). We also observed very limited growth in broth culture when *MSMEG_0916* was induced with ATc (**Fig. S6**). Similar experiments done in *Mycobacterium bovis* BCG (BCG) and MTB with Rv0472c overexpression, also resulted in 3-log (**Fig. 5B**) and 4-log (**Fig. S7**) reduction in viability, respectively.

**Fig. 5.**
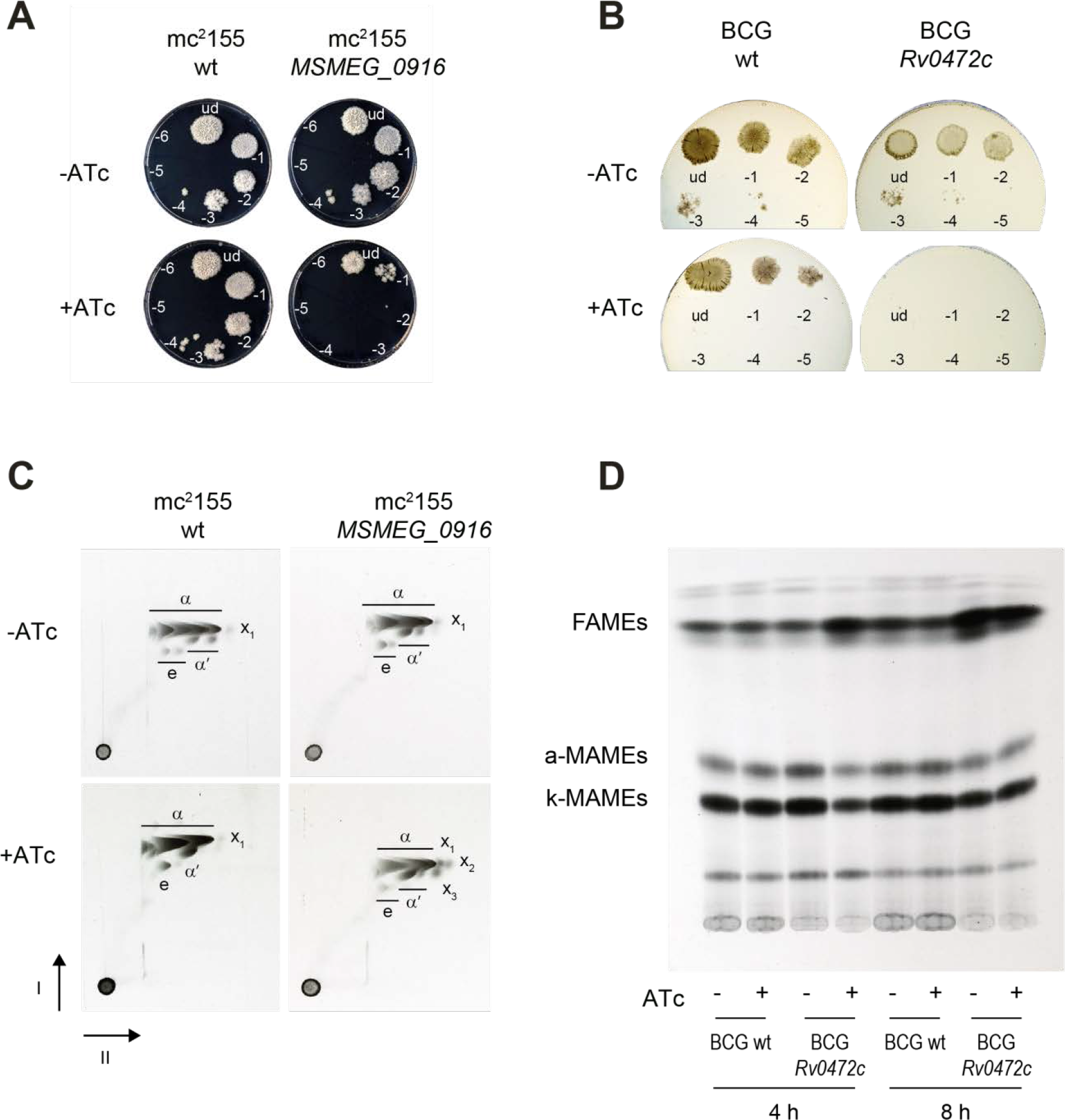
Cell viability and mycolic acid characterization. (**A**) Serial ten-fold dilutions of MSM wild type (wt) and MSM with inducible overexpression of *MSMEG_0916* were spotted on 7H10 agar plates with or without ATc. (**B**) Serial ten-fold dilutions of BCG wt and BCG with inducible overexpression of *Rv0472c* were spotted on 7H10 agar plates with or without ATc. (**C**) Argentation TLC of ^14^C-labeled methyl esters of mycolic acids (MAMEs) obtained from apolar lipids and delipidated cell wall fractions of MSM wt and MSM with inducible overexpression of *MSMEG*_0916. The α, α′, epoxy (e) and cyclopropanated α- (X_1_) MAMES species are labeled. Faster-migrating species that co-migrated with α-MAMES and accumulate with induced *MSMEG*_0916 overexpression are indicated as X_2_ and X_3_. (**D**) BCG wt and BCG with inducible overexpression of *Rv0472c* cultures, labeled with ^14^C-acetate, were induced (+ATc) or uninduced (−ATc) for 4 h or 8 h. The total FAMEs and MAMEs were extracted and analyzed by autoradiography-TLC using equal counts (15,000 cpm) for each lane.

Overexpression of the conserved TF resulted in a loss of mycobacterial viability, due to repression of *desA1* and *desA2* and ensuing decrease in mycolic acid biosynthesis. Conditional depletion of DesA1 in MSM leads to an intermediate decrease in desaturation prior to complete loss of mycolic acids(*17*). To test for a decrease in mycolic acid biosynthesis, we labeled cultures of *MSMEG_0916* overexpression strain with ^14^C acetic acid following growth in the presence or absence of ATc. Thin layer chromatography (TLC) analysis of apolar lipids demonstrated that overexpression of *MSMEG_0916* reduced the levels of trehalose dimycolates (TDMs) (**Fig. S8**). We also analyzed methyl esters of mycolic acids (MAMEs) obtained from apolar lipids using 2D-argentation TLC analysis, designed to separate each subclass of mycolic acid based on saturation levels. MAMEs analysis revealed an accumulation of products that migrate identically to our previous observation with DesA1 depletion(*17*) (**Fig. 5C**) and most likely correspond to mono-unsaturated mycolates. Similarly, 1D TLC separation of fatty acid methyl esters (FAMEs) and MAMEs from apolar lipids confirmed the general decrease of MAMEs and an accumulation of FAMEs when Rv0472c is overexpressed in BCG (**Fig. 5D** and densitometric analysis in **Fig. S9**). This characteristic profile of total MAMEs inhibition and FAMEs accumulation mirrors what is seen with fatty acid synthase (FAS)-II inhibitors, such as isoniazid(*43*) and thiolactomycin(*44*), and confirms the involvement of DesA1 and DesA2 in the biosynthesis of mycolic acids and more specifically with the FAS-II system. Interestingly, in BCG there was no accumulation of mono-unsaturated mycolates as those found in MSM upon detailed analysis of MAMEs by 2D-argentation TLC (**Fig. S10A** and **S10B**). This could be due to key differences in mycolate subclasses between MSM and MTB, particularly cyclopropane ring formation, which is abundant in MTB but not MSM and requires a precursory desaturation event.

## CONCLUSIONS

During MTB infection, the bacterium utilizes various mechanisms to ensure its own survival and persistence in the host. Mycolates, essential for mycobacterial cell wall rigidity and stability, have been prime candidates for such virulence factors(*45*). Mycolic acids not only make up a lipid rich barrier in the mycobacterial cell envelope, they also act as potent immunomodulators, driving the pathogenesis of MTB, primarily as part of the cord factor (TDM)(*46*, *47*). Here, we present evidence that mycolate biosynthesis is tightly regulated in response to the intracellular environment. Using our novel Path-seq method, we observed significantly induced expression of the fatty acid desaturases, *desA1* and *desA2*, 24 h after MTB infection of mice. We further demonstrated that *desA1* and *desA2* are regulated by Rv0472c (*MSMEG*_0916) and that Rv0472c-mediated repression leads to reduced mycolate biosynthesis and loss of mycobacterial viability. As DesA1 and DesA2 have been shown to be involved in mycolic acid biosynthesis via desaturation of the merochain, we have therefore named their transcriptional regulatory protein MadR (for <u>M</u>ycolic <u>a</u>cid <u>d</u>esaturase <u>R</u>egulator).

Not much is known about the regulation of mycolic acid biosynthesis apart from two transcription factors shown to regulate distinct operons, both containing genes encoding core FAS-II proteins(*19*, *48*). Studying the regulatory alterations to mycolate subclasses remains an even greater challenge, especially during infection. Our studies show that MadR is involved in the *in vivo* and *in vitro* regulation of *desA1* and *desA2*, aerobic desaturases involved in mycolate merochain desaturation(*17*). The introduction of double bonds in the merochain precedes cyclopropanation and other merochain modifications that are critical for pathogenic mycobacteria(*49*–*52*). Loss of cyclopropanation can lead to hyperinflammatory responses and attenuated infection. As the introduction of double bonds in the merochain is required for subsequent cyclopropanation and other merochain modifications, DesA1 and DesA2 could be drivers of both mycolic acid biosynthesis and composition during infection. In other words, MadR driven regulation not only leads to lower mycolate levels during dormancy, a state when new cell wall material is not synthesized, but also altered cyclopropane ring formation by varying desaturation levels, thus affecting virulence and persistence.

Surprisingly, we observed early (2 h post infection) induced expression of *desA1* and *desA2* during MTB infection of BMDMs, followed by return to basal levels by 8 h. This is consistent with the reported increased production of TDM within the first 30 min after *in vitro* phagocytosis(*53*) and suggests that the desaturases play a role in cell wall modifications that occur in response to intracellular cues. However, the presence of these cues appears to be different or delayed in AMs from infected mice. The overall disparity in the transcriptional profile of MTB from BMDMs and AMs is both intriguing and disturbing. The active MTB networks we identified from BMDMs imply the presence of early and sustained bacterial stress. However, the induction of these stress-related genes is absent in the transcriptomes of MTB from AMs, suggesting the bacteria are not experiencing the same type or amount of stimuli in AMs. These data support recent observations using fluorescent MTB reporter strains, demonstrating that bacilli in AMs exhibit lower stress and higher bacterial replication than those in interstitial macrophages(*24*). Similarly, we hypothesize that MTB responds divergently to macrophages of different lineages and that AMs present fewer stresses and possibly a more permissive environment compared to BMDMs. It is also worth mentioning that the data raises some concerns with respect to the use of BMDMs as an appropriate infection model.

The expression dynamics of *desA1* and *desA2* during MTB infection of BMDMs is also mirrored in RNA-seq data of MTB entering and exiting hypoxia over a 5 day time course (**Fig. 6A**). In this experiment, we used mass flow controllers to regulate the amount of air and nitrogen (N_2_) gas streaming into cultures of MTB and achieve a gradual depletion of oxygen over 2 days (**Fig. 6A**). The cultures were maintained in hypoxia for 2 days by streaming only N_2_ then reaerated over 1 day by a controlled increase in air flow. During the 2 day oxygen depletion, the expression levels of *desA1* and *desA2* did not change significantly. However, as soon as the cultures reached hypoxia, the expression of the desaturases increased for ~5h, followed by a dramatic repression after ~30h of being in hypoxia. Subsequently, reaeration of the culture returned *desA1* and *desA2* to basal expression levels (**Fig. 6B**). This data along with the results described above leads us to propose a model for MadR regulation of *desA1* and *desA2* transcription as summarized in **Fig. 6C**. Under normal growth conditions, MadR exists in equilibrium between the free and DNA-bound forms, thus, maintaining basal levels of *desA1* and *desA2* transcripts. Upon macrophage infection and early hypoxia, equilibrium favors unbound MadR which de-represses *desA1* and *desA2* transcription and increases mRNA levels. As infection progresses and reaches later stages of hypoxia, MadR has increased binding affinity in the promoters of *desA1* and *desA2* and represses their transcription to below basal levels. Ultimately, the MadR regulatory system enables mycobacteria to efficiently alter mycolate biosynthesis and composition in response to environmental signals. We suspect the early response to infection (*desA1* and *desA2* up-regulation) increases desaturation events and allows MTB to fine-tune cyclopropanation and other merochain modifications that contribute to the establishment of infection. However, mycolate biosynthesis is energetically expensive and MadR-mediated repression occurs in later stages of infection. The reduction in mycolate biosynthesis allows MTB to enter dormancy and facilitates long term persistence.

**Fig. 6.**
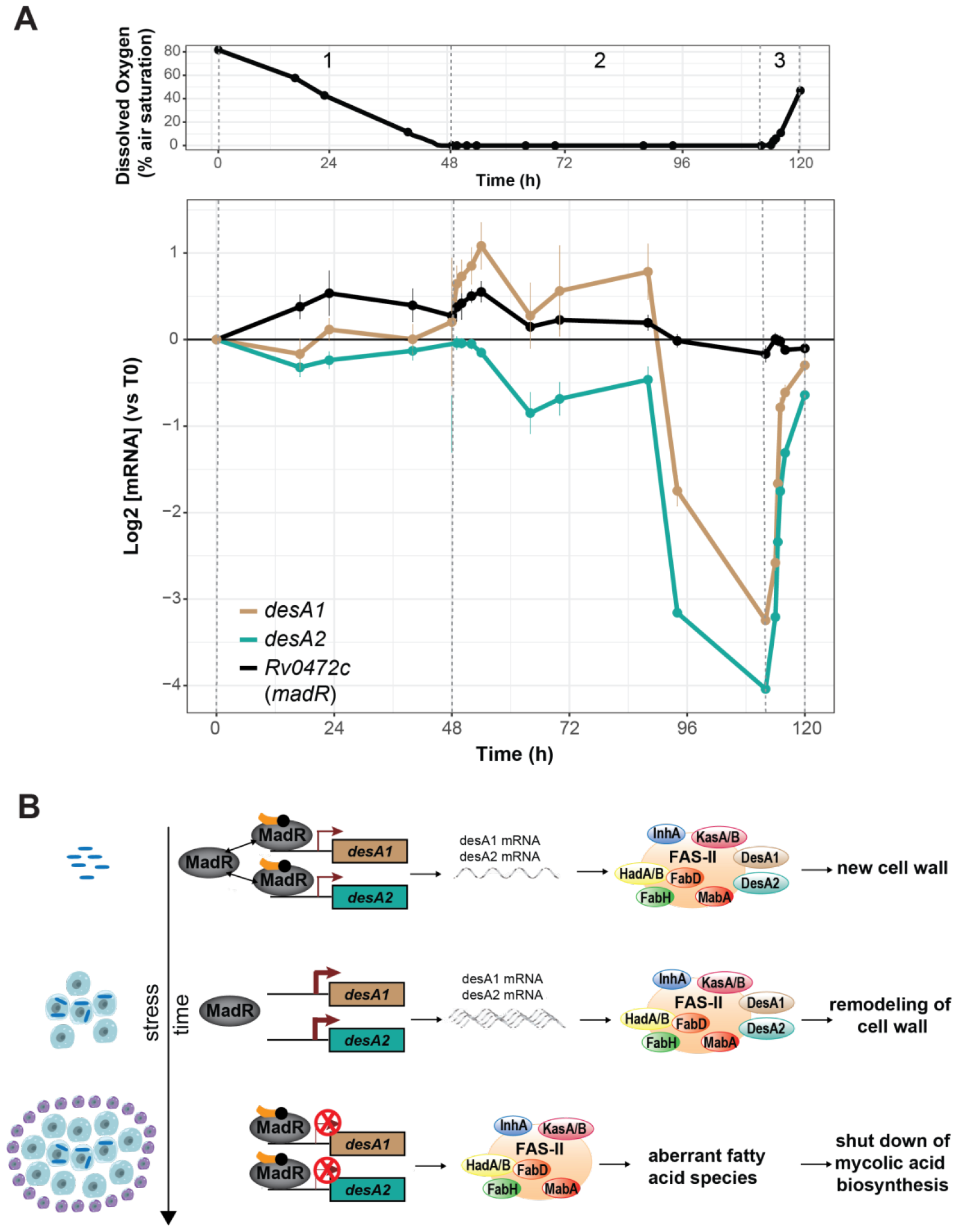
Expression of *desA1*, *desA2* and *Rv0472c* during hypoxia time course and model of MadR regulation of desaturases. (**A**) Dissolved oxygen curve over 120 h time course showing controlled depletion (*1*), sustained hypoxia (*2*) and controlled reaeration (*3*). Points represent the average of three biological replicates and were measured via fiber optic technology that non-invasively probes oxygen levels in the culture (PreSens Precision Sensing GmbH). (**B**) Expression profiles (RNA-seq) of *desA1*, *desA2* and *Rv0472c (madR)* over the time course and oxygen levels. Error bars show the standard deviation from three biological samples. (**C**) Under normal growth conditions, MadR exists in equilibrium between the free and DNA-bound forms, and a basal level of *desA1* and *desA2* expression is maintained. DesA1 and DesA2 acts with other enzymes of the fatty acid synthase-II (FAS-II) complex to produce mycolic acids for new cell wall. Infection of macrophages and early stress cues shifts the equilibrium towards the free form of MadR and the repression of *desA1* and *desA2* is released. The desaturase protein levels increase, introducing double bonds that allow cyclopropanation and other modifications to alter the cell wall. These changes enable the bacteria to withstand intracellular stresses and establish infection. As infection continues and stress is sustained, MadR binds tightly to the promoter of desA1 and desA2, leading to stringent repression of the desaturases. MadR-mediated repression of desA1 and desA2 leads to irregular fatty acids of medium length and the pausing of mycolic acid biosynthesis to enter into dormancy. The DNA-bound form of MadR is shown in complex with a yet unknown co-factor that leads to repression of the desaturases.

The question remains how MadR is able to differentially bind to DNA in response to environmental changes. In mycobacteria, the other TFs regulating mycolate biosynthesis are modulated by long chain acyl-CoAs(*54–56*), proposing a role for these molecules in the modulation of MadR as well. Similarly, a MadR homolog in *Pseudomonas aeruginosa*, DesT, was shown to have enhanced DNA binding in the presence of unsaturated acyl-CoAs(*57*). These studies support the notion that a select acyl-CoA ligand may control MadR DNA binding affinity (as shown in **Fig. 6C**), and thus the expression of *desA1* and *desA2*.

The characterization of the MadR regulon provides valuable insight for understanding the evolution of MTB. While we have shown the regulation by MadR is conserved from MSM to MTB, our results also suggest the fatty acid desaturation events and resulting mycolate subclasses have evolved, specializing for bacterial survival in the host environment. These findings propose mycobacterial evolution from saprophyte to pathogen has occurred through the adaptation of ancestral genes and regulatory networks to function in the host environment. Ultimately, this study demonstrates the *in vivo* significance of the desaturases and their regulation by MadR. We believe the Path-seq method, described and employed here, offers a sensitive and tractable approach to elucidate the molecular mechanisms used by MTB during host infection. Our detailed characterization of one such mechanism has revealed that modulation of MadR activity can affect mycolate composition as well as mycobacterial viability. Accordingly, we have established Path-seq as a powerful tool for uncovering the minimally studied *in vivo* biology of this pathogen and revealed the essentiality of MadR encoded program for cell wall remodeling and biosynthesis. As such, we present MadR as a new and important anti-tubercular target.

## Acknowledgements

We thank members of the Baliga, Bhatt and Aderem labs for critical discussions; Albel Singh for help with graphics; David Sherman and lab for generating the overexpression system and MTB Rv0472c overexpression data.

## Author contributions

E.J.R.P, R.B., A.C.R., A.B., and N.S.B. designed research; E.J.R.P, R.B., A.C.R., A.K., M.P., D.M., and C.C. performed research; E.J.R.P and M.A performed computational analyses; E.J.R.P, R.B., A.C.R., A.B., and N.S.B. analyzed data; and E.J.R.P, R.B., A.B., and N.S.B. wrote the paper.

## Data and materials availability

Sequencing data have been deposited in the GEO repository in a SuperSeries record under accession number GSE116085. Correspondence and requests for materials should be addressed to NSB.

## Materials and Methods

The approaches used in this study include both computational and biological methods. Algorithms developed for the EGRIN model and the NetSurgeon approach were implemented in the R programming language and Python, respectively. Plots were generated using R and images prepared using Adobe Illustrator CS5.

### Culturing conditions

Mycobacteria strains were cultured in Middlebrook 7H9 with the ADC supplement (Difco), 0.05% Tween80 at 37°C under aerobic conditions with constant agitation. Strains containing the anhydrotetracycline (ATc)-inducible expression vector were grown with the addition of 50 μg/mL hygromycin B to maintain the plasmid. Growth was monitored by OD600 and colony forming units (CFUs). For experiments featuring *madR* overexpression strains, overexpression was induced for the approximate duration of one cell doubling (18h for MTB and BCG, 4h for MSM) using an ATc concentration 100 ng/mL culture. Wild-type and overexpression strain cultures were grown into mid-log phase. For assessing growth on agar plates, broth cultures were adjusted to OD600 of 0.5, and serial dilutions were spotted on 7H10 containing 0.5% (v/v) glycerol, and 10% (v/v) OADC plates, with or without 100 ng/ml ATc. In the case of the overexpression strain, 50 ng/ml hygromycin was added to the solid medium. For growth in broth, MSM mid-logarithmic phase cultures containing the integrative vector pMV306-eGFP-Zeo were inoculated in an initial OD600 0.05 in 200 μl of 7H9 supplement with 0.2% (v/v) glycerol, 0.05% (v/v) Tween-80 and 10% (v/v) OADC with or without 100 ng/ml ATc in sealed 96-well Costar 3603 black sided clear bottomed plate incubated at 37°C. Fluorescence was acquired every 30 min for 30 hours at 37°C, in a Pherastar FS microtiter plate reader (BMG Labtech), using 485 nm and 520 nm as excitation and emission wavelengths, respectively. When growing BCG or MSM for lipid extraction, cultures were cultured up to OD600 0.5. Then, they were induced with 100 ng/ml ATc (Sigma-Aldrich) final concentration, and labelled with acetic acid [1-14C] 1 mCi/ml (PerkinElmer), if hot lipid analysis was going to be performed. MSM samples were collected the following day, while BCG cultures at 4 and 8 hours post-induction.

### Tissue Culture

BMDMs were cultured in RPMI (RPMI containing 10% (v/v) FBS, 2mM L-glutamine) with recombinant human CSF-1 (50 ng/mL) for 6 days and then replated. BMDMs were infected on day 7 with MTB H37Rv strain (MOI 10), followed by washing 3x with RPMI at 2 h post infection. MTB infected BMDMs were lysed with TRIzol (Invitrogen) and total RNA was isolated from mixed host-pathogen sample.

### Strains

To investigate the growth properties of MadR overexpression, we used strains containing an ATc-inducible expression vector of the gene, as described previously(20, 26, 27, 58). The pDTCF-*MSMEG_0916* plasmid was transformed into *M. smegmatis* mc^2^155. Smilarly, pDTCF-*Rv0472c* was transformed into *M. bovis* BCG and *M. tuberculosis* H37Rv. The same *M. tuberculosis* H37Rv was used for Path-seq experiments.

### Mice

C57BL/6 mice were purchased from the Jackson Laboratory. All mice were housed and bred under specific pathogen-free conditions at the Center for Infectious Disease Research (CID Research). All experimental protocols involving animals were approved by the Institutional Animal Care and Use Committee of CID Research.

### Aerosol Infection

A mid log-phase stock of MTB H37Rv was used to infect mice in an aerosol infection chamber (Glas-Col). Bacterial load in the lungs was determined by plating serial dilutions from homogenized lungs.

### Cell Isolation, Analysis and Sorting

Bronchoalveolar lavage was performed by first exposing the trachea of euthanized mice. The exposed trachea was punctured using Vannas Micro Scissors (VWR) and 1 mL PBS was injected using a 20G-1” IV catheter (McKesson) connected to a 1 mL syringe. The PBS was flushed into the lung and aspirated three times and the recovered fluid was placed in a 15mL tube on ice. The 1 mL PBS wash was then repeated 3 additional times for a total of 4 mL recovered fluid. Cells were filtered, spun down and resuspended in a 96-well plate for antibody staining. Cells were suspended in 1X PBS (pH 7.4) containing 0.01% NaN_3_ and 1% fetal bovine serum (i.e. FACS buffer). Fc receptors were blocked with anti-CD16/32 (2.4G2, BD Pharmingen). Cell viability was assessed using Zombie Violet dye (Biolegend). Surface staining included antibodies specific for murine Siglec F (E50-2440, BD Pharmingen), CD11b (M1/70, Biolegend), CD64 (X54-5/7.1, Biolegend), CD45 (104, Biolegend), CD3 (17A2, eBiosciences), and CD19 (1D3, eBiosciences). Cell sorting was performed on a FACS Aria (BD Biosciences). Cells were collected in complete media, spun down, resuspended in Trizol, and frozen at - 80° overnight prior to RNA isolation.

### RNA isolation

Cell pellets in TRIzol were transferred to a tube containing Lysing Matrix B (QBiogene, Inc.), and vigorously shaken at max speed for 30 s in a FastPrep 120 homogenizer (QBiogene) three times. This mixture was centrifuged at max speed for 1 min and the supernatant was transferred to a fresh tube. RNA from extracellular MTB samples, BMDM infection and madR overexpression was isolated using the Direct-zol RNA MicroPrep kit (Zymol Research) according to manufacturer’s instruction with on-column DNase treatment. RNA from mice infection was isolated by adding 200 μL chloroform. Samples were inverted and incubated for 2-3 min and the upper aqueous phase was collected. A second cholorform extraction was done, followed by addition of 1 μL glycolgen and 500 μL isopropanol. Samples were incubated with isopropanol for 10m at room temperature, centrifuged and supernatant was discarded. Pellet was washed with 1 ml 70% ethanol twice. All ethanol was removed, the pellet dried (15m) and resuspended in 12 μL RNase free water. Total RNA yield was quantified by Nanodrop (Thermo Scientific) and quality was analyzed in a 2100 Bioanalyzer system (Agilent Technologies). Total RNA samples were depleted of ribosomal RNA using the Ribo-Zero Gold rRNA Removal Kit “epidemiology” (Illumina).

### Probe design

Nonoverlapping head-to-tail 120-nucleotide probes were designed using the Array software (Agilent Technologies). A total of 35,624 probes were designed to cover 3,924 *M. tuberculosis* H37Rv ORFs (assembly M_tub_h37rv_ASM19595v2_32_1). Using Megablast, it was verified that all genes of MTB were matched by at least one probe and that only a negligible fraction of the probes could be mapped on the mouse and human cDNA sequences from Ensembl.

### Preparation of libraries for transcriptional sequencing

RNA libraries for Path-seq were prepared using the SureSelect^XT^ strand-specific RNA target enrichment for Illumina multiplexed sequencing. RNA libraries for RNA-seq were prepared using the SureSelect^XT^ strand-specific RNA kit, but were not hybridized to probes. Briefly, mRNA was enzymatically fragmented and double-stranded cDNA was produced with adapters ligated to both ends. The library was then amplified using provided primers which hybridize to the previously inserted adapters, therefore allowing a linear amplification to all transcripts present in the sample. In the case of nonenriched RNA-seq samples, sample indexes were also inserted during this PCR. For Path-seq libraries, double-stranded cDNA ligated to adapters was also amplified and then incubated at 65 Cfor 24h with the set of biotinylated oligonucleotides specifically desgined to capture MTB transcripts, as described above. The hybridized sequences were captures with magnetic streptavidin beads. They were next linearly amplified using provided primers and indexed during PCR. Before sequencing, libraries were assessed for quality and fragment size by Bioanalyzer and with a Qubit fluorometer (Invitrogen) to determine cDNA concentration. Resulting libraries were sequenced on the Illumina NextSeq instrument using mid output 150 v2 reagents. Paired-end 75 bp reads were processed following Illumina default quality filtering steps.

### Transcription abundance from sequencing data

Raw FASTQ read data were processed using the R package DuffyNGS as described previously(*59*). Briefly, raw reads pass through a 3-stage alignment pipeline: (i) a prealignment stage to filter out unwanted transcripts, such as rRNA, mitochondrial RNA, albumin, and globin; (ii) a main genomic alignment stage against the genome(s) of interest; and (iii) a splice junction alignment stage against an index of standard and alternative exon splice junctions. Reads from samples of mixed host-pathogen RNA and extracellular MTB controls were aligned to a combined *M. tuberculosis H37Rv* (ASM19595v2) and *Mus Musculus* (GRCm38.p6) genome. Reads from samples of MSM RNA were aligned to *M. smegmatis* mc^2^155 genome (ASM1500v1). All alignments were performed with Bowtie2(*60*), using the command line option “very-sensitive.” BAM files from stages 2 and 3 are combined into read depth wiggle tracks that record both uniquely mapped and multiply mapped reads to each of the forward and reverse strands of the genome(s) at single-nucleotide resolution. Multiply mapped reads are prorated over all highest-quality aligned locations. Gene transcript abundance is then measured by summing total reads landing inside annotated gene boundaries, expressed as both RPKM and raw read counts. Two stringencies of gene abundance are provided using all aligned reads and by just counting uniquely aligned reads.

### Differential expression

For both infection models (*in vitro* and *in vivo*), we used DESeq2(*61*) to identify gene expression changes between intracellular and extracellular MTB at each sampled time point. We used rounded raw read counts estimated by DuffyNGS (as described above) as input for DESeq2. Genes with absolute log2 fold change bigger than one and multiple hypothesis adjusted *P*-value below 0.01 and 0.05, for the in vitro and in vivo data, respectively, were considered differentially expressed.

For *MSMEG_0916* overexpression, we used a panel of 5 DE tools to identify gene expression changes between induced (+ATc) and uninduced (−ATc). The tools included (i) RoundRobin (in-house); (ii) RankProduct(*62*); (iii) significance analysis of microarrays (SAM)(*63*); (iv) EdgeR(*64*); and (v) DESeq2(*61*). Each DE tool was called with appropriate default parameters and operated on the same set of transcription results, using RPKM abundance units for RoundRobin, RankProduct, and SAM and raw read count abundance units for DESeq2 and EdgeR. All 5 DE results were then synthesized, by combining gene DE rank positions across all 5 DE tools. Specifically, a gene’s rank position in all 5 results was averaged, using a generalized mean to the 1/2 power, to yield the gene’s final net rank position. Each DE tool’s explicit measurements of differential expression (fold change) and significance (*P*-value) were similarly combined via appropriate averaging (arithmetic and geometric mean, respectively). Genes with averaged absolute log2 fold change bigger than one and multiple hypothesis adjusted *P*-value below 0.01 were considered differentially expressed.

### MTB signed transcriptional network

We compiled a signed (stating the positive or negative nature of each TF-gene interaction) wiring diagram of MTB transcriptional regulatory network. The compiled MTB network included 4,635 TF-gene interactions (2,296 and 2,339 instances of activation and repression, respectively) with both physical (detected with ChIP-seq experiments) and functional evidence (detected with transcriptional profiling). The compiled network contained 1,996 genes and 136 TFs with at least one target. The initial ChIP-seq derived MTB network consisted of 6,581 interactions occurring in the −150bp to +70bp region of genes’ promoter reported by Minch et al.(*26*). We expanded that MTB ChIP-seq network by taking into account operon organizations. For a given TF-gene interaction, if the target gene is part of an operon (according to the MicrobesOnline database(*65*)), we included all other members of the operon as potential targets of the corresponding TF. The expanded MTB ChIP-seq network contained 12,188 interactions. Finally, we filtered out interactions that did not change at least 20% in the relevant TF-over-expressing strain (compared to the WT strain). Up-regulation of the target gene in the TF-over-expressing strain was interpreted as positive interaction (the opposite for down-regulation).

### Identification of transcription factors with differential activity in intracellular MTB

We applied the strategy recently proposed by Michael *et al.*(*25*) (the NetSurgeon algorithm) to identify potential TFs with an increase or decrease in their regulatory activity in intracellular MTB with respect to the extracellular controls at each sampled time point. Briefly, we used a signed MTB transcriptional network model (described above) and the lists of genes identified as differentially expressed by DESeq2 and their computed adjusted *P*-values (described above), to score and rank the contribution of TFs’ activity to the observed changes in the transcriptional profiles of intracellular MTB. To reduce false positives (i.e. misleading presence of TFs with small number of known targets at the top of our TF scores ranking) due to overlap between regulons, only TFs with at least five targets were considered in this analysis. Furthermore, we considered TFs in the top 15 of NetSurgeon’s score ranking as the ones with differential activity. We selected this threshold based on empirical analysis in *Escherichia coli*.

### ChIP-seq

The madR overexpression strain was induced for the approximate duration of one cell doubling (4 h for MSM) using an ATc concentration of 100 ng/ml culture. DNA-protein interactions were characterized as described previously(*26*). Libraries were prepared using ThruPLEX DNA-seq Kit (Rubicon) using standard protocol. Samples were sequenced on the Illumina 550 NextSeq instrument, generating unpaired 20-30 million 75-bp reads per sample. Raw FASTQ read data were processed using the R package DuffyNGS as described previously(*59*). For consensus motif determination, we searched for conserved DNA sequences within ±50 nucleotides of high quality (score > 0.7) ChIP-seq peak centers using MEME(*66*).

### Measuring viability of *madR* overexpression strains

Wild-type and *madR* overexpression strain cultures were grown into mid-log phase. For assessing growth on agar plates, OD of the broth culture was adjusted up to 0.5, and serial dilutions were spotted in 7H10 containing 0.5% (v/v) glycerol, and 10% (v/v) OADC plates, with or without 100 ng/ml ATc. In the case of the overexpression strain, 50 ng/ml hygromycin was added to the solid medium.

### Extraction and analysis of total lipids and mycolic acids

Lipids were extracted from BCG and MSM cells in three fractions as describes by Dobson *et al.*(*67*), with a few modifications. Briefly, outside apolar lipids from dried pellets were extracted with two consecutive extractions with 4 ml of petroleum ether (60-80C), and dried. Then, inside apolar and polar lipids were extracted following Dobson protocol.

Outside, inside apolar and polar lipid extracts, along with delipidated pellets from MSM and BCG were subjected to alkaline hydrolysis using tetrabutylammonium hydroxide (TBAH) as previously described(*68*).

Aliquots (15,000 cpm) from each outside, inside apolar and polar lipid extracts were analyzed by thin layer chromatography (TLC) utilizing Silica Gel 60 F254 plates (Merck) developed once in the solvent system CHCl_3_/CH_3_OH/H_2_O (60:16:2, v/v/v). However, FAMEs and MAMEs aliquots (15,000 cpm) were resolved through TLC using petroleum ether/acetone (95:5, v/v) or by two-dimensional silver ion argentation thin layer chromatography (2D-TLC)(*68*). Autoradiograms were produced after exposing Carestream^®^ Kodak^®^ BioMax^®^ MR film for 3 days. To determine the intensity of TLC spots, densitrometric analysis using Adobe Photoshop CC 2015 was performed.

**Fig. S1.**
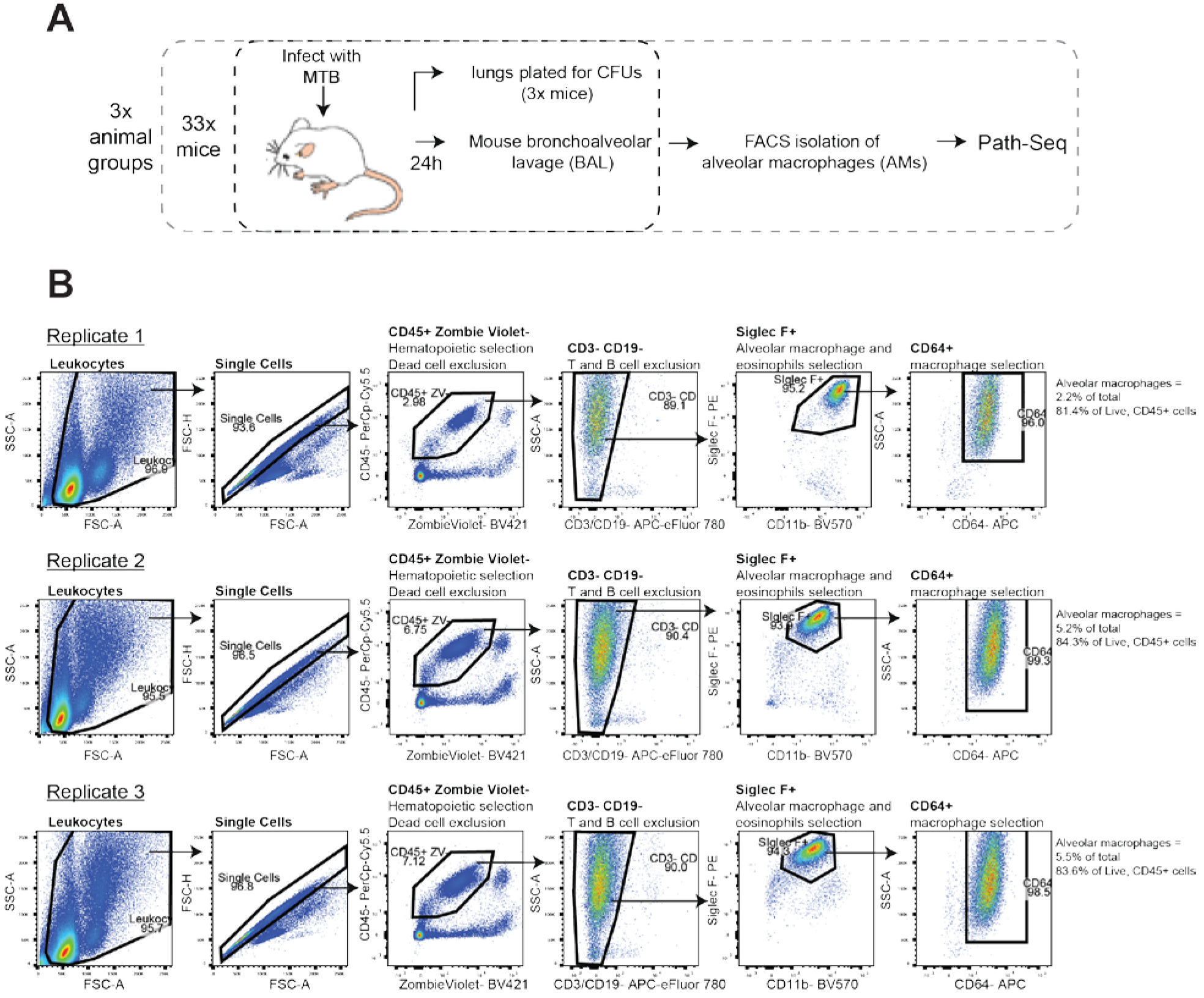
Isolation of alveolar macrophages from MTB infected mice. (**A**) Schematic of *in vivo* infection using Path-seq method. (**B**) Flow cytometry analysis to sort alveolar macrophages from BAL of wt mice 24 h after aerosol infection of 6×10^3^ MTB. Cell viability was assessed using Zombie Violet viability dye. Alveolar macrophages were defined as CD45^+^, CD3^−^, CD19^−^, SiglecF^+^, CD11b^mid^ and CD64^+^. Plots show the percentages of cells from each parent population for each of the three infection replicates.

**Fig. S2.**
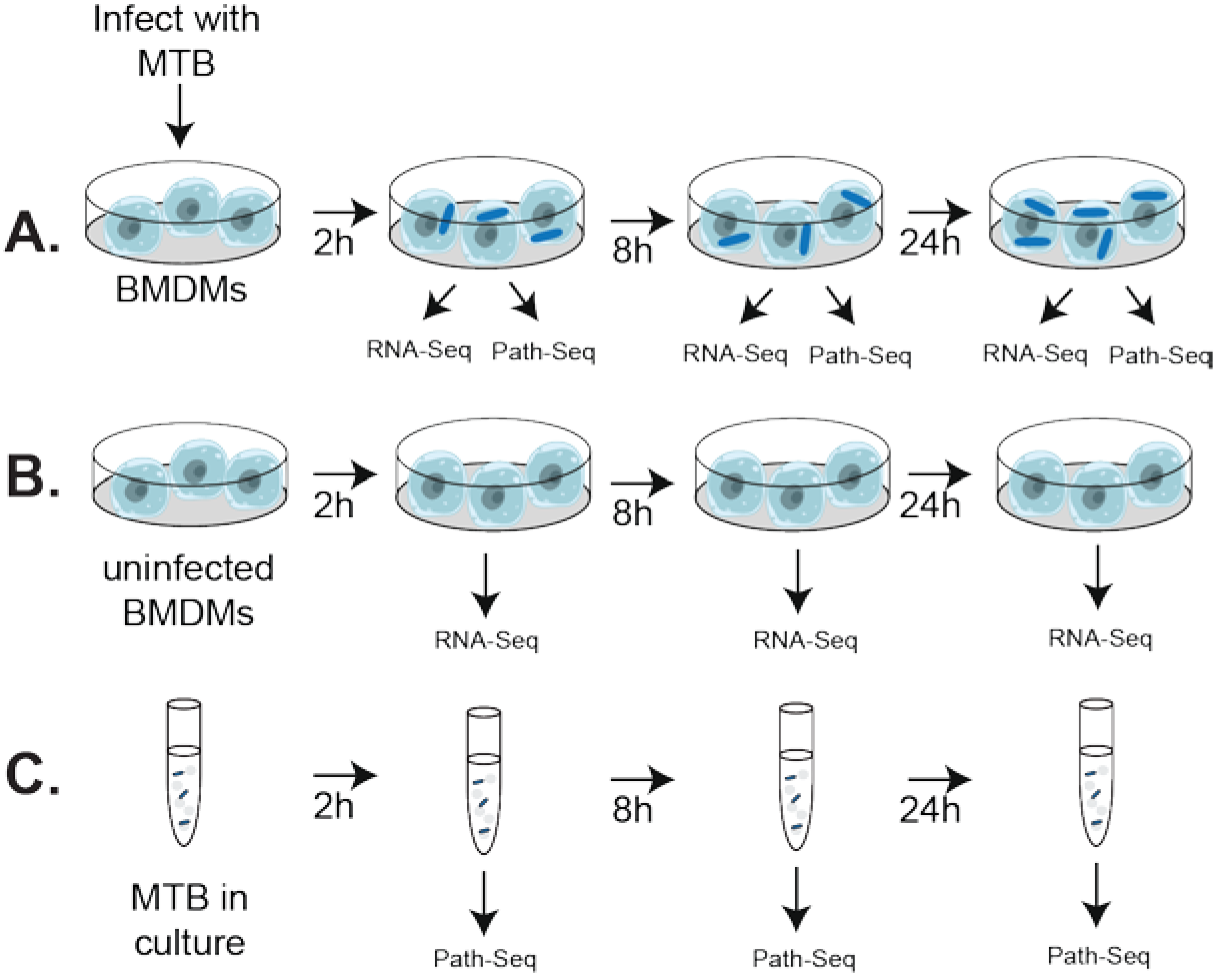
Schematic of *in vitro* infection using Path-seq. (**A**) Mouse bone marrow derived macrophages (BMDMs) were infected with MTB H37Rv at MOI of 10. MTB infected BMDMs were lysed with TRIzol at given time points and RNA samples were prepared for sequencing by RNA-seq and Path-seq (MTB enrichment), as described in main text. (**B**) Uninfected BMDMs were collected as a host control and processed by RNA-seq. (C) MTB grown in 7H9 broth culture were used as extracellular MTB control and processed by Path-seq.

**Fig. S3.**
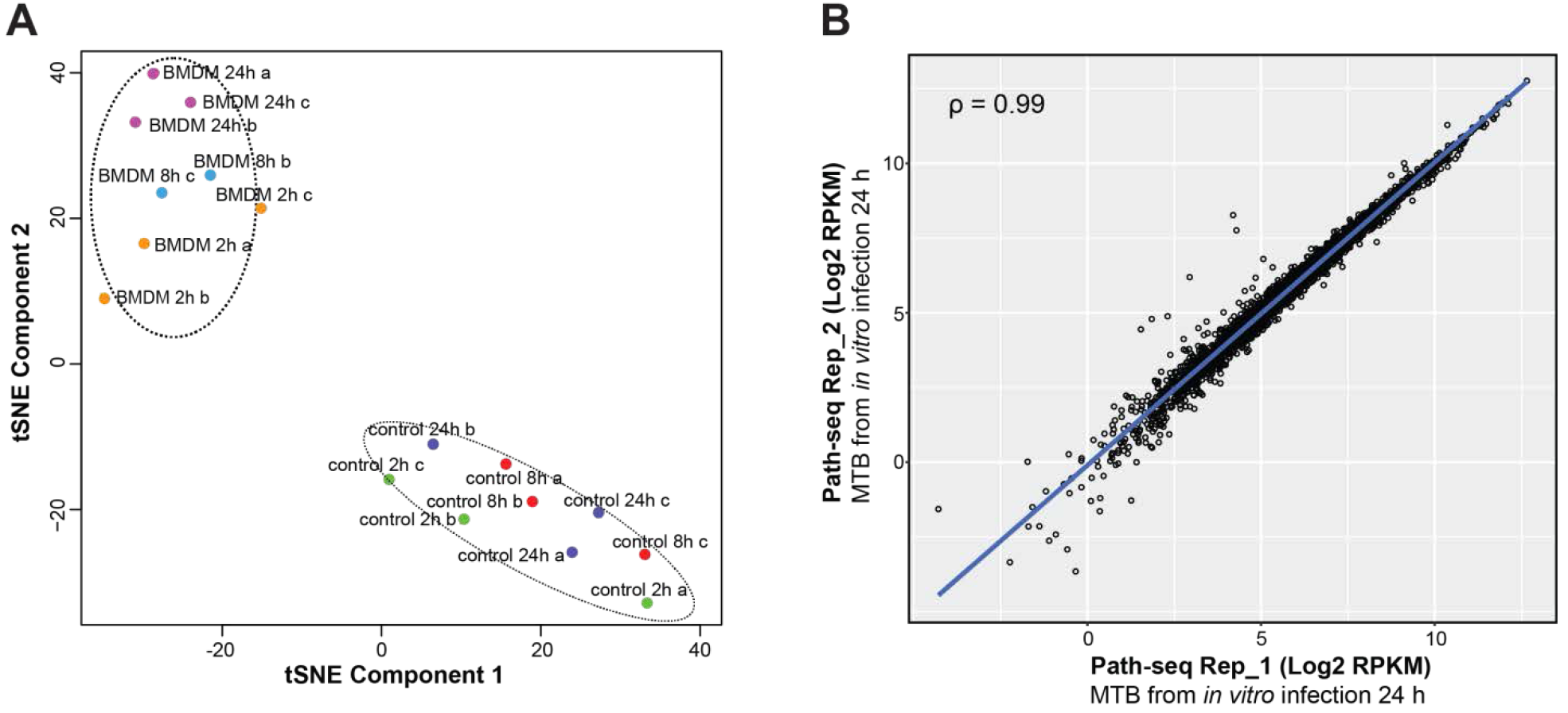
Analysis of replicates from *in vitro* infection. (**A**) tSNE analysis of Path-seq data from *in vitro* infection samples. MTB from infected bone marrow derived macrophages are labeled as “BMDM” and extracellular MTB grown in 7H9 are labeled as “control”. (**B**) Correlation between replicates from *in vitro* infection collected at 24 h. Scatter plot of log2 RPKM values is shown with Pearson correlation, *P*-value < 0.0001.

**Fig. S4.**
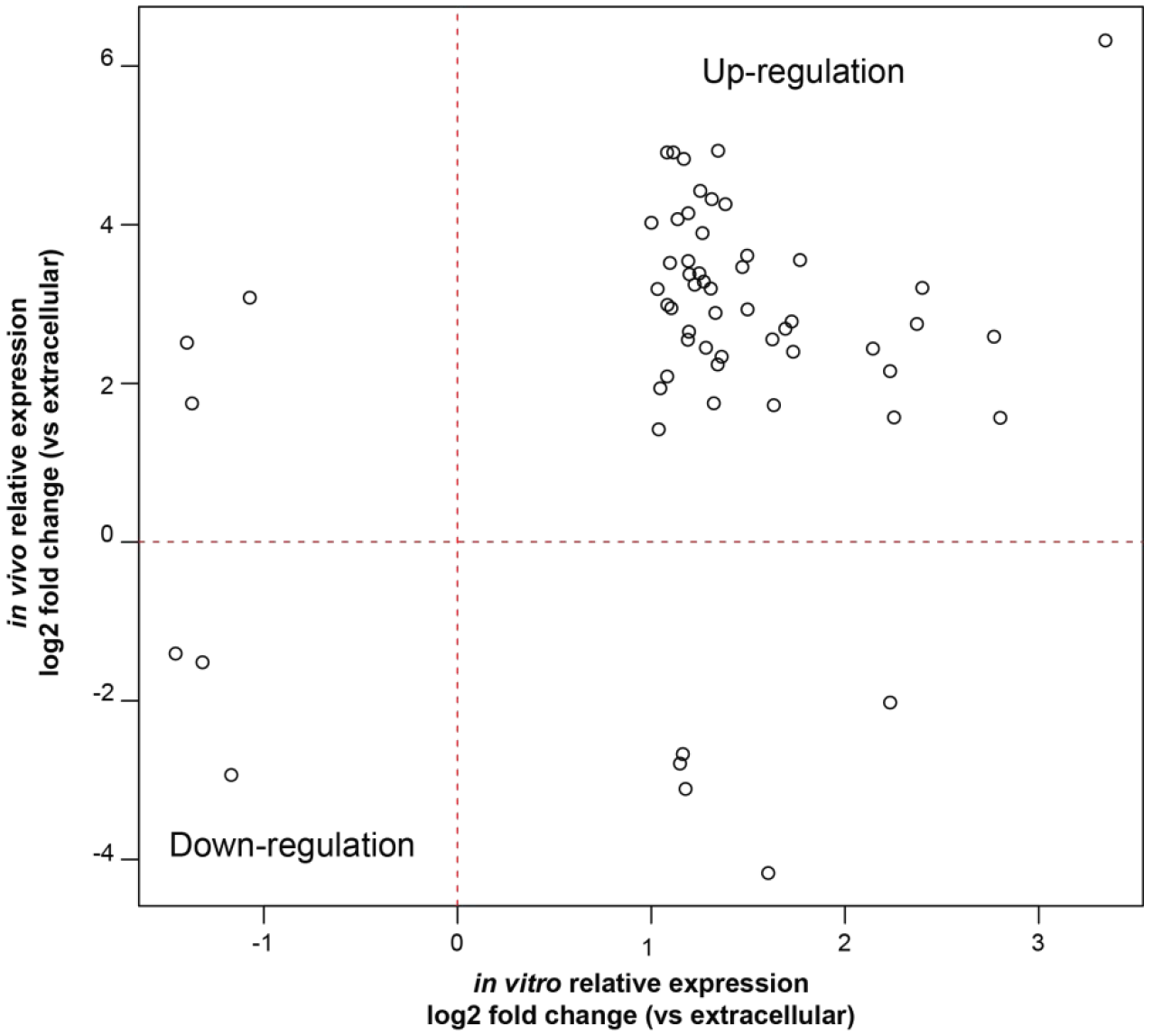
Expression of genes that are significantly differentially expressed from both in vitro and in vivo infection models. Scatter plot of log2 fold change expression of MTB from infected BMDMs vs extracellular MTB at 24 h (in vitro) and log2 fold change of MTB from alveolar macrophages of MTB infected mice vs extracellular MTB at 24 h. All samples were processed by Path-seq method.

**Fig. S5.**
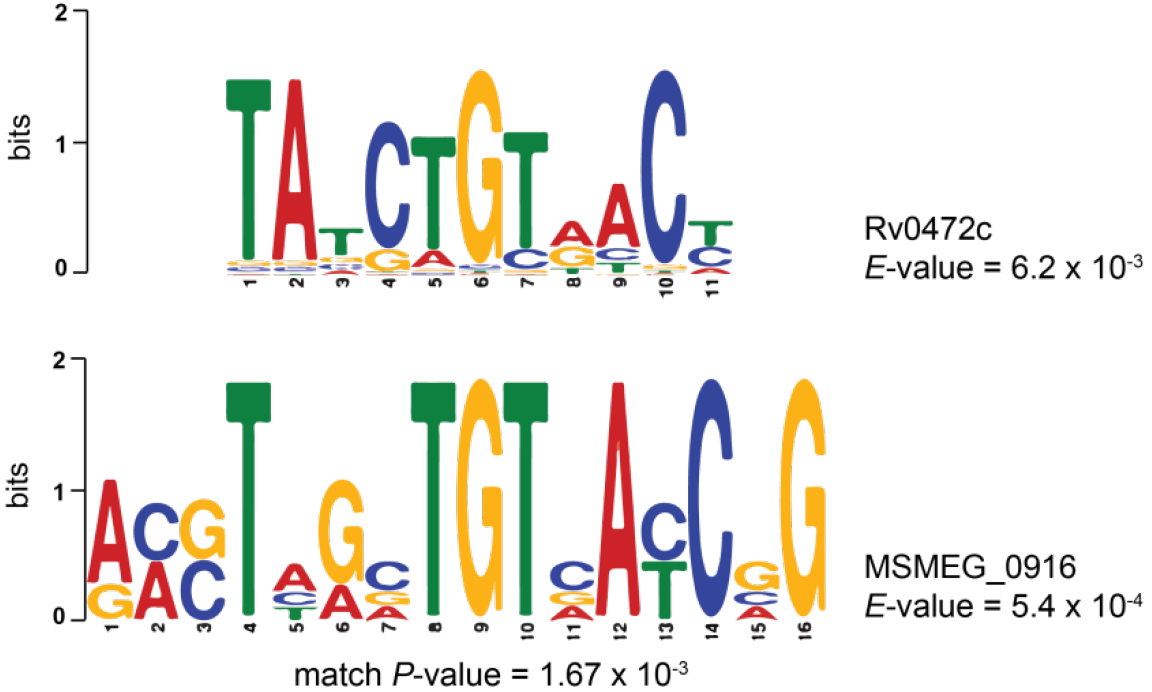
Consensus motifs from ChIP-seq peaks of *Rv0472c* overexpressed in MTB (top) and *MSMEG_0916* overexpressed in MSM (bottom). For consensus motif determination, we searched conserved DNA sequences within ±50 nucleotides of high quality (score > 0.7) ChIP-seq peak centers using MEME(*66*). Alignment of consensus motifs was performed with Tomtom(*69*), match *P*-value = 1.67×10^−3^.

**Fig. S6.**
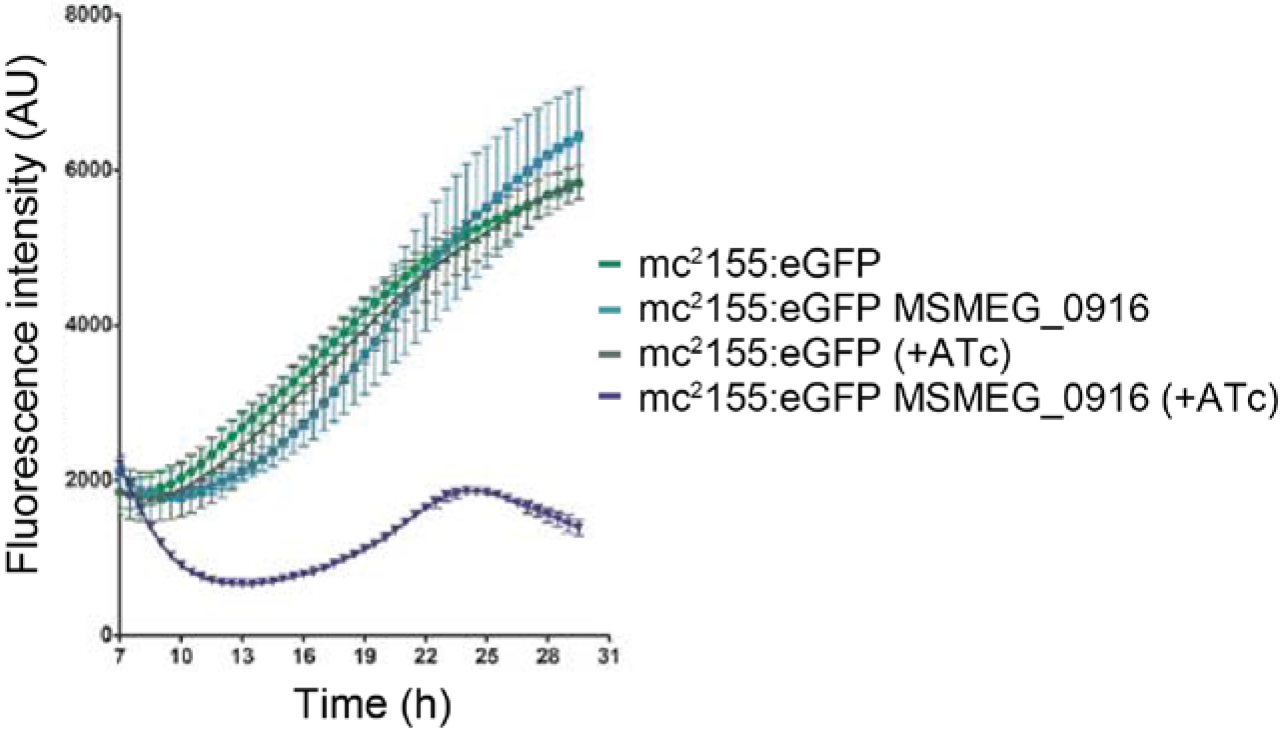
Growth of MSM overexpressing *MSMEG_0916* in 7H9 broth. MSM mc^2^155 wt and mc^2^155/pDTCF-MSMEG_0916 strains were transformed with an eGFP integrative vector. Growth of mc^2^155 wt and mc^2^155/pDTCF-MSMEG_0916 was monitored for fluorescence intenstiy (485/520 nm) in 96-well plates in the presence or absence of ATc.

**Fig. S7.**
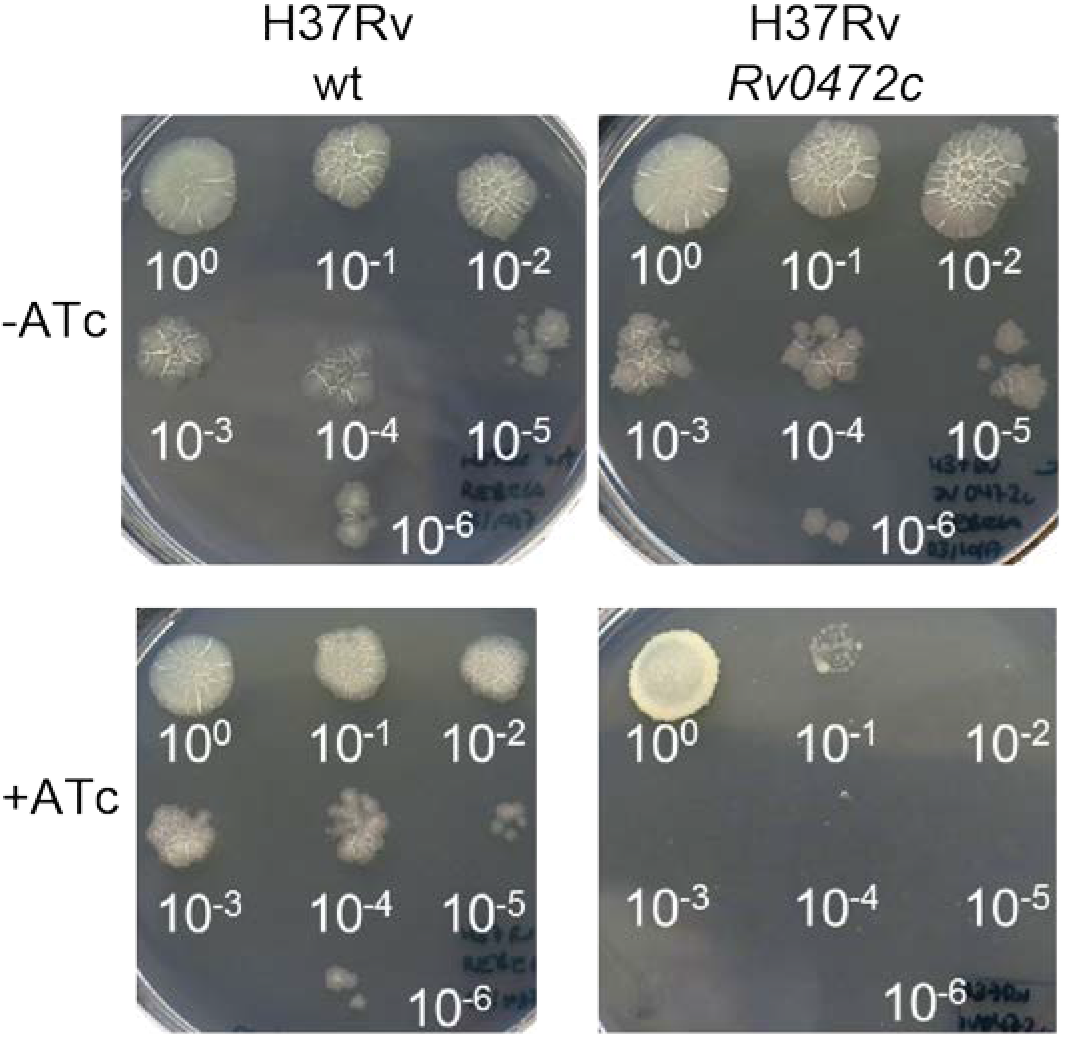
Cell viability of MTB overexpressing *Rv0472c*. Serial ten-fold dilutions of MTB H37Rv wild type (wt) and MTB with inducible overexpression of *Rv0472c* were spotted on 7H10 agar plates with or without ATc.

**Fig. S8.**
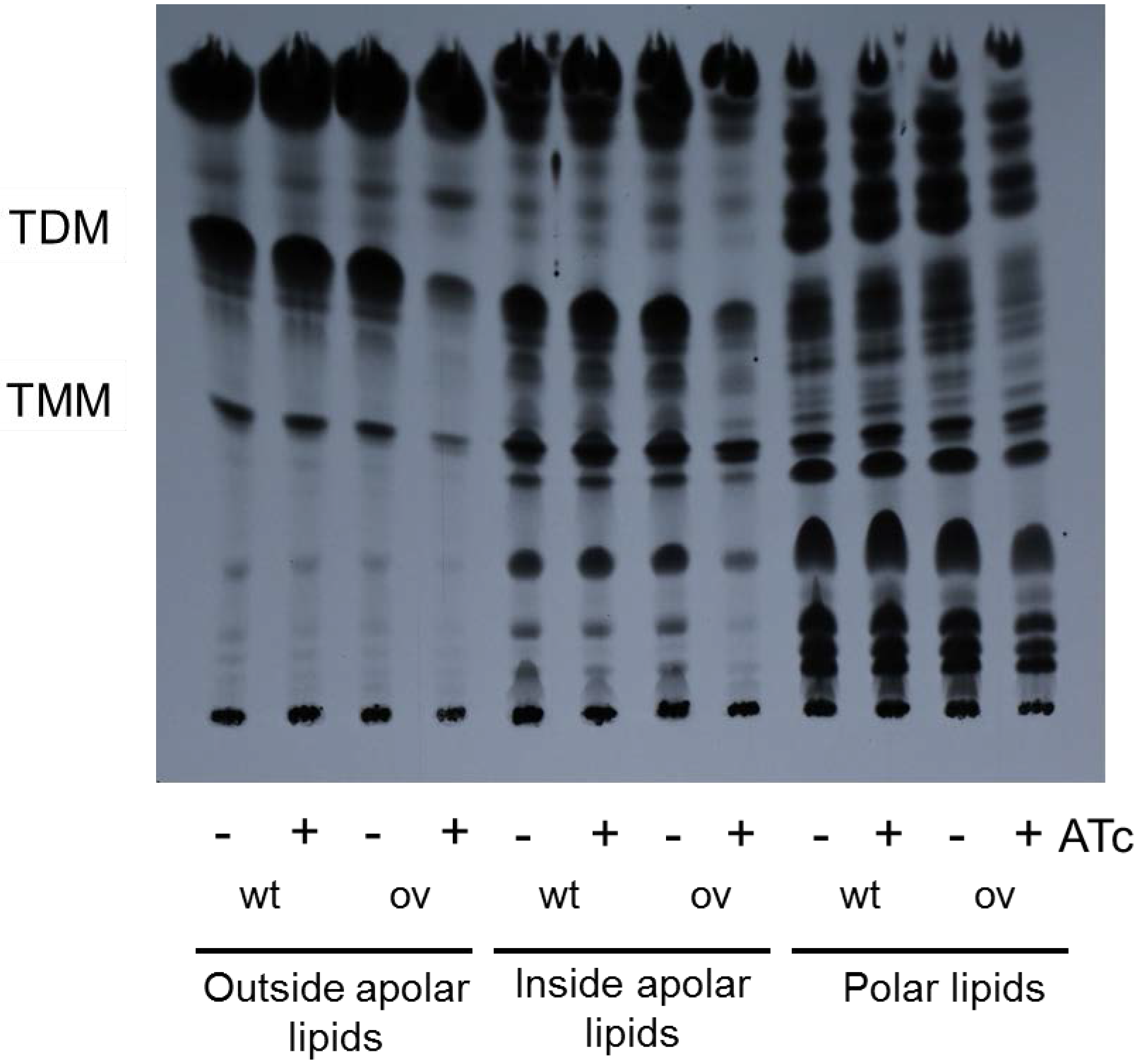
TLC of outside, inside and apolar lipids extracted from MSM wildtype (wt) and MSM overexpressing *MSMEG_0916* (ov) in the presence (+) or absence (−) of ATc. Trehalose dimycolate (TDM) and trehalose monomycolate (TMM) were extracted following labeling and analysed by autoradiography-TLC using equal counts (15,000 cpm) for each lane.

**Fig. S9.**
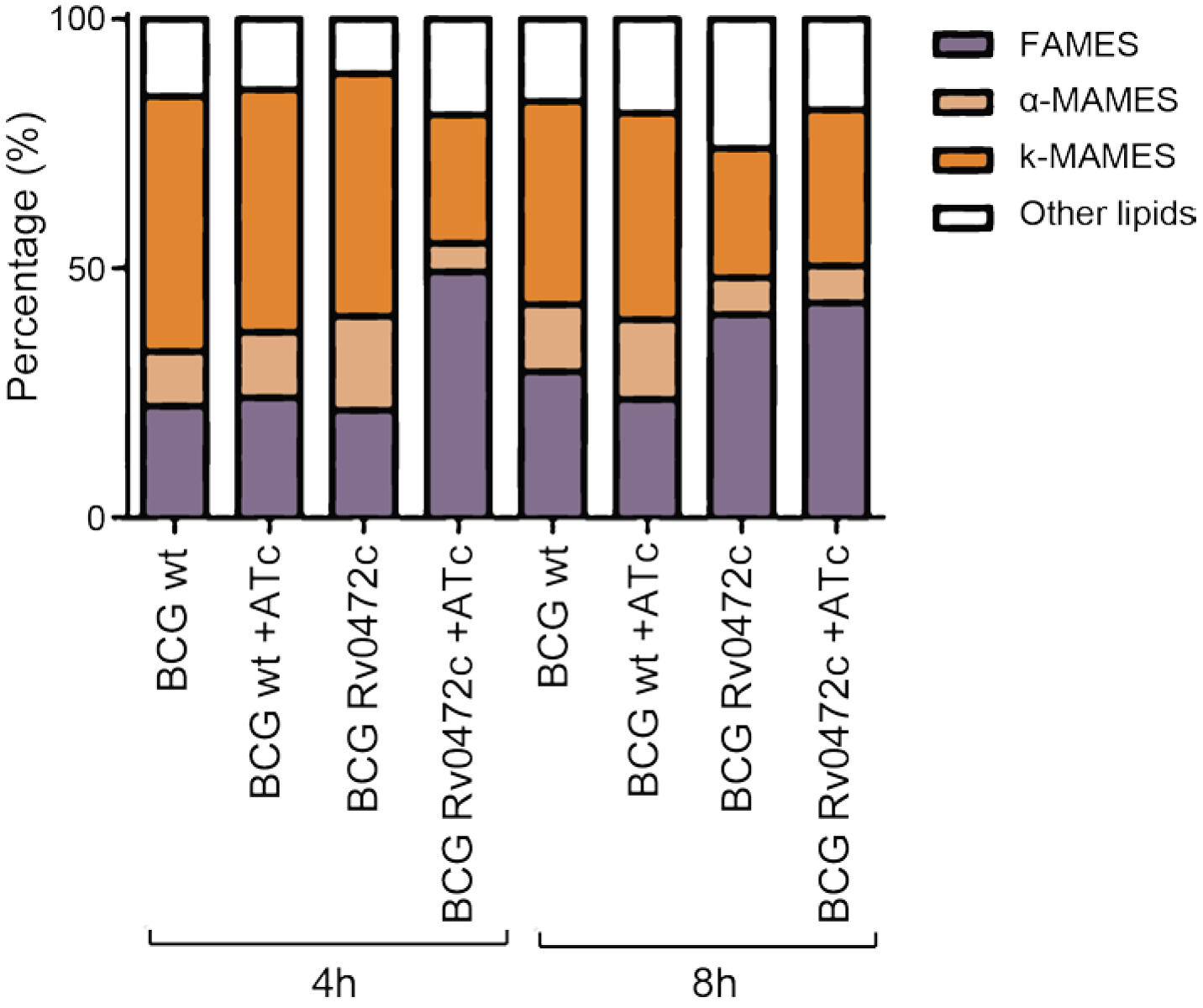
Bar graph showing the relative amounts of ^14^C-labeled methyl esters from BCG wildtype (wt) and BCG overexpressing *Rv0472c* species with or without ATc for 4 h and 8 h. The methyl ester amounts are indicated as percentages of total amounts of ^14^C-labeled methyl esters detected on the TLC plate shown in **Fig. 5D**, as determined by densitometry. FAMES; fatty acyl methyl esters. α-MAMES; α-mycolic acid methyl esters. k-MAMES; keto-mycolic acid methyl esters.

**Fig. S10.**
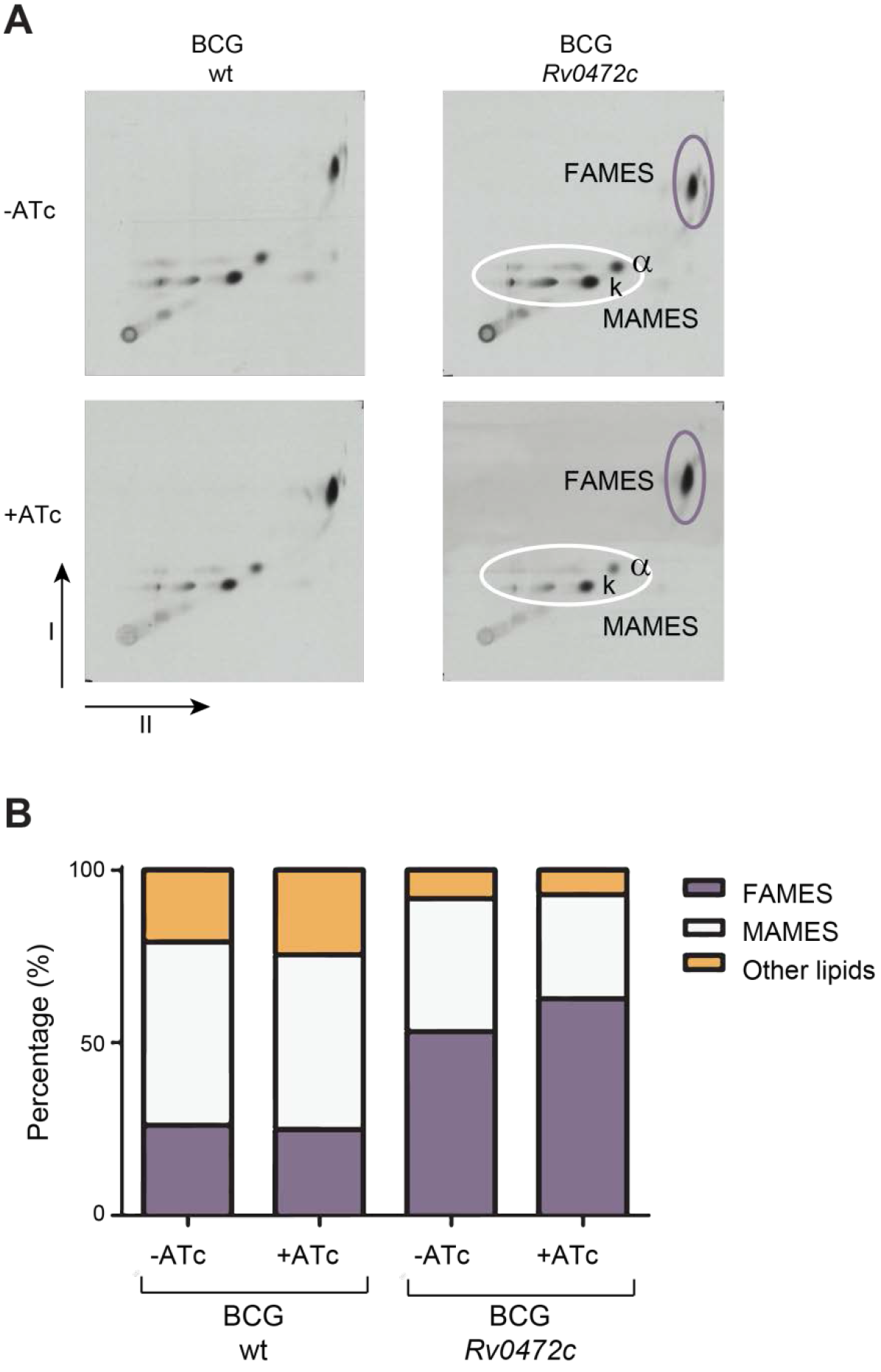
TLC analysis of FAMEs and MAMEs from BCG overexpressing *Rv0472c*. (**A**) 2D-argentation TLC analysis of fatty acid methyl esters (FAMEs) and mycolic acid methyl esters (MAMEs) from inside apolar lipid extracts of BCG wildtype (wt) and BCG overexpressing Rv0472c in the presence or absence of ATc. Alpha(α)-MAME and keto(k)-MAME species are indicated. (**B**) Bar graph showing the relative amounts of ^14^C-labeled MAMES and FAMES as percentages of total amounts of ^14^C-labeled methyl esters detected on the 2D TLC plates shown in **A**, as determined by densitometry.

**Table S1.** Summary analysis from Path-seq data of alveolar macrophages (AMs) from MTB infected mice and extracellular MTB grown in 7H9 broth.

|  | Non-zero genes (%) | Total read counts | Mean count | Noise |
| --- | --- | --- | --- | --- |
| AM T24h (deposition = 8100 CFUs) | 49.88 | 997861 | 354 | 1.95 |
| AM T24h (deposition = 5633 CFUs) | 31.23 | 1077327 | 611 | 2.03 |
| AM T24h (deposition = 4333 CFUs) | 14.4 | 183380 | 226 | 1.81 |
| extracellular 24h | 98.22 | 1588332 | 286 | 2.01 |
| extracellular 24h | 98.32 | 2867574 | 517 | 1.96 |
| extracellular 24h | 97.85 | 1603959 | 290 | 1.99 |

### Data S1. (separate file)

Significantly differentially expressed genes of MTB from *in vivo* infection vs extracellular MTB using Path-seq.

### Data S2. (separate file)

Significantly differentially expressed genes of MTB from *in vitro* infection vs extracellular MTB using Path-seq.

